# Thriving Across Depths: How Blue Light Shapes a Large PSI Supercomplex and Specific Photosynthetic Traits in the seagrass *Posidonia oceanica*

**DOI:** 10.1101/2025.06.20.660723

**Authors:** Quentin Charras Ferroussier, Charlie Mathiot, Dmitry A. Semchonok, Eduard Elias, Ahmad Farhan Bhatti, Régine Lebrun, Dorian Guillemain, Marina I. Siponen, Roberta Croce, Colette Jungas

## Abstract

Photosynthetic organisms rely on finely tuned mechanisms to optimize photosynthesis under different light conditions. While these processes are well-characterized in land plants, the adaptive strategies of marine plants remain largely unexplored. The Mediterranean seagrass *Posidonia oceanica* (Alismatales), a key ecosystem engineer thriving from the surface up to 40m depth and one of the largest long-term blue carbon sinks in coastal environments. Here, we investigate how *P. oceanica* adjusts its photosynthetic apparatus in response to varying light spectra encountered at different seawater depths. Contrary to land plants, *P. oceanica* maintains a relatively high PSI/PSII ratio and a high content of the major light-harvesting complex II (LHCII), regardless of depth. Notably, the antenna size of the photosystems remains stable across depths, although we document significant depth-dependent reorganization of the thylakoid membrane ultrastructure. Moreover, we identify a novel large PSI-LHCII supercomplex (L-PSI-LHCII) in *P. oceanica*, characterized by additional Lhca proteins, reduced red-shifted absorption, and increased chlorophyll *b* content. Ultrafast spectroscopy reveals the distinct energy transfer dynamics within this complex. The presence of a similar supercomplex in other marine plants, such as *Zostera marina*, suggests a conserved adaptive strategy among seagrasses.

## Introduction

**Blue carbon** refers to carbon sequestered by vegetated coastal ecosystems, including **salt marshes**, **mangroves**, and **seagrass meadows** over millennia. Although seagrasses occupy only 0.01% of the Earth’s surface, they account for approximately 4% of global carbon stocks (Duarte et al., 2013). Among these, *Posidonia oceanica*, a Mediterranean endemic, forms extensive meadows from shallow waters down to 40 m depth (Pasqualini et al., 1998), covering roughly 25% of the region’s coastal seabeds (Telesca et al., 2015). Its slow horizontal expansion (1–5 cm/year) enables the formation of persistent structures, with some meadows estimated to be up to 80,000 years old (Arnaud-Haond et al., 2012).

*P. oceanica* is a key ecosystem engineer supporting forming ecological niches for a multitude of taxa (Díaz-Almela and Duarte, 2008; Boudouresque et al., 2012) and is recognized as a highly efficient carbon sink. The reported long-term sequestration of ∼270 t C/ha/year (Pergent-Martini et al., 2021), significantly surpassing the ∼4 t C/ha/year sequestration typical of tropical forests (Yadav et al., 2022). This sequestration capacity corresponds to 11–42% of historical COL emissions from Mediterranean countries since the Industrial Revolution (Pergent et al., 2014), primarily through the matte, the dense, decay-resistant layer of rhizomes, roots and sediment (Boudouresque et al., 2016).

Despite its ecological and biogeochemical significance, *P. oceanica* meadows have experienced a decline of approximately 34% over the past five decades (Pergent et al., 2014) with current annual losses exceeding 1.5%. Anthropogenic pressures including coastal development, increased turbidity, mechanical damage from anchoring, and rising sea surface temperatures are the primary drivers of decline (Marbà et al., 1996; Boudouresque et al., 2012). While restoration initiatives, such as transplantation, have yielded variable outcomes (Molenaar, 1992; BOUDOURESQUE et al., 2013), a more comprehensive understanding of the physiological and biochemical mechanisms underlying *P. oceanica*’s adaptation and resilience is urgently needed for conservation and management strategies under accelerating global change.

In marine environments, both light intensity and quality rapidly decline with depth, with red light disappearing entirely beyond ∼10 m and overall intensity reduced 20 to 50 times at deeper meadows (see below). *P. oceanica* thrives across these varying conditions and understanding its adaptive mechanisms provides valuable insights into photosynthetic processes. Photosystems I (PSI) and photosystem II (PSII) and their antenna systems (LHC) drive linear electron flow to produce ATP and NADPH, used for downstream COL fixation. Optimal electron transfer requires balanced excitation between PSI and PSII, maintained through acclimation strategies such as adjusting PSI/PSII ratios (Chow et al., 1990; Wientjes et al., 2017), LHCII content (Wagner et al., 2008; Kouřil et al., 2013; Albanese et al., 2016), antenna size, and state transitions (Goldschmidt-Clermont and Bassi, 2015; Grieco et al., 2015; Rantala et al., 2020). Underwater, the dominant blue-green wavelengths are primarily absorbed by chlorophyll *b*, abundant in LHCII linked mostly to PSII (Su et al., 2017), enhancing its absorption capacity compared to PSI-LHCI (Bos et al., 2023). Thus, PSI excitation may become limited with increasing depth.

Most of our knowledge about photosynthetic regulation mechanisms come from land model plants grown under artificial light conditions. However, how photosynthesis is regulated at the molecular level in the plant’s natural habitats remains poorly documented. The present study adresses this underexplored area. Here, we examined the depth-dependent photosynthetic adaptations of *P. oceanica*, focusing on LHCII levels, PSI/PSII stoichiometry, and antenna sizes at depths of 2, 15, and 26 m. We unexpectedly found stable, elevated LHCII content and PSI/PSII ratios across all depths compared to land angioserpm. Using a new protocol developed to isolate thylakoids from polyphenol-rich marine plants (Charras et al., 2024), we identified a Chl *b*-enriched PSI supercomplex, consisting of canonical PSI-LHCI linked to phosphorylated LHCII trimers and an additional Lhca1-Lhca4 heterodimer, uniformly expressed at all tested depths. Notably, the far-red absorbing Chls typical of land plants (Morosinotto et al., 2003; Romero et al., 2009) were absent in PSI of *P. oceanica* and related Mediterranean marine species, highlighting a common blue-shifted adaptation in all tested seagrasses, and the expression of the L-PSI-LHCII in deep-growing marine plants.

## Results

### The light environment of the *P. oceanica* meadows selected for the study

The light spectrum at each depth tested was determined (**Suplemental Figure 1**). From the surface to a depth of 2 m, light intensity decreases by a factor of ∼2.5 (from 2000 to 800 µmol.m^−2^.s^−1^) largely in the 600-750 nm region. However, at depths of 15 and 26 m, the irradiance is between 20 and 50 times lower than at the surface (120 and 40 µmol.m^−2^.s^−1^, respectively) and completely depleted of red light; only green and blue-green light remain, peaking around 450-500 nm.

**Figure 1:**
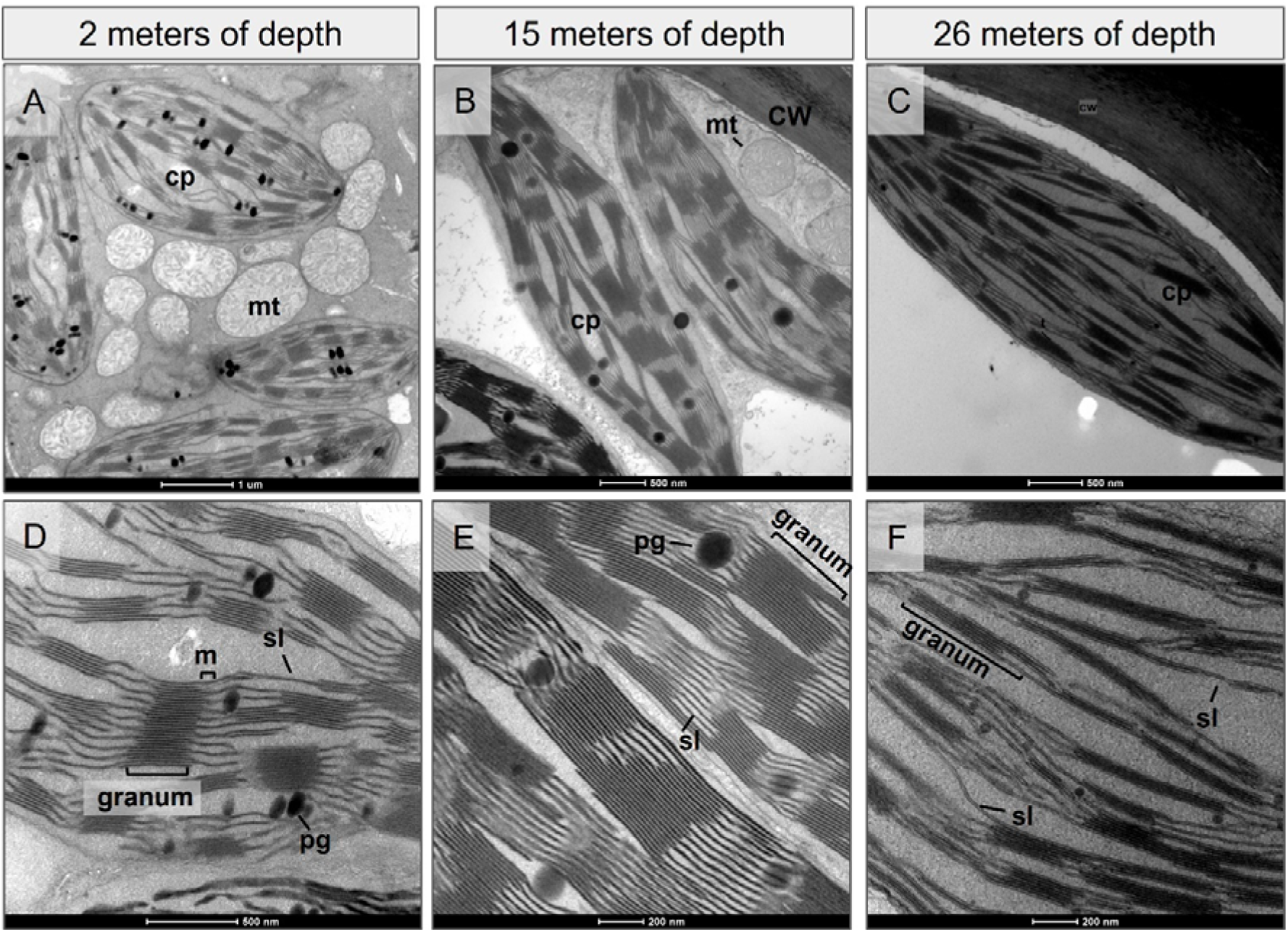
Transmission electron microscopy (TEM) analysis of the *P. oceanica* chloroplasts from three growing depths. Electron micrographs of negative-stained chloroplasts cross section (up) and thylakoid membrane (down) from *P. oceanic*a leaves that grew at **(A-B)** 2 m depth, **(C-D)** 15 m depth, and **(E-F)** 26 m depth. **cp**: chloroplast, **cw**: cell wall, **mt**: mitochondria, **m**: margins, **pg**: plastoglobules, **sl**: stroma lamellae. The corresponding scale bars are specified below each figure panel.

### Depth-dependent changes in chloroplast ultrastructure

Long-term acclimation of plants to varying light conditons occurs through the composition and structural organization of the thylakoid membrane (Buschmann et al., 1978; Ferroni et al., 2016; Schumann et al., 2017). The chloroplast acclimation of *P. oceanica* was examined at the structural level in leaves collected at different depths (2, 15, and 26 m) using electron microscopy (**Figure 1**). The micrographs and the grana size quantification (**Suplemental Figure 2**) revealed significant alterations in membrane organization: At 15 m, the grana thickness increased **(Figure 1B and 1E)** compared to 2 m (200 nm vs. 140 nm) (**Figure 1A and 1D**), while the grana diameter remained unchanged (around 420 nm). Surprisingly, at 26 m (**Figure 1C and 1F**), grana thickness decreased (80 nm vs. 140 nm at 15 m), but their diameter was larger (510 nm vs. 420 nm at 15 m). Similar observations have been made in *Arabidopsis thaliana* exposed to light favoring PSII excitation (Wagner et al., 2008), consistent with the predominance of blue-green light in deeper water.

**Figure 2:**
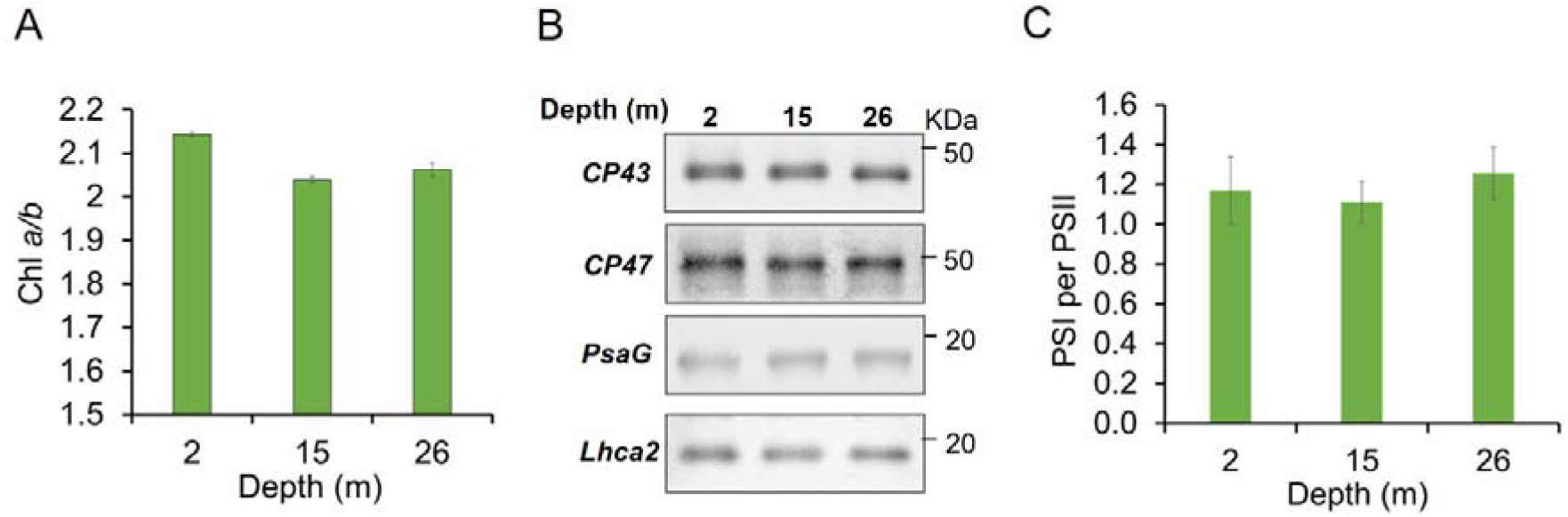
Chlorophyll content and photosystems stoichiometry in *P. oceanica* thylakoids at different growing depths. (**A**) Chl *a/b* of the thylakoid membranes of *P. oceanica* collected at 2, 15, and 26 m depth in winter. Data for each depth are expressed as the mean ± SD of 2 technical replicates from a thylakoid preparation obtained with 30 shoots (4 leaves/shoot). **(B)** Immunoblot of thylakoid membranes from *P. oceanica* collected at 2, 15, and 26 m depth. 0,5 μg of Chl was loaded on the gel. The primary antibodies used are indicated. Quantification of the band’s intensity is reported in Suplemental Figure 4a. **(C)** PSI/PSII stoichiometry ± SD in fresh young winter leaves estimated by difference of ECS signal between untreated (PSII+PSI) and DCMU-HA infiltrated (PSI only) leaves. Data are expressed as the mean from two independent experiments (**Suplemental Figure 4D and 4E**) at the 3 tested depths (n=3 different leaves from each depth and for each experiment).

### *P. oceanica* shows a low Chl a/b ratio at all depths

Changes in the Chl*a/b* ratio is a proxy of changes in the number of LHCII. Surprisingly, in *P. oceanica*, we found that the Ch*la/b* ratio varied little at the depths tested (2 m: 2.14±0.004, 15 m: 2.04±0.007, 26 m: 2.06±0.02) (**Figure 2A**). This ratio is very low compared with that of *A. thaliana*, which is around 3.4-3.7 in high light and 2.7-2.9 in low light (Kouřil et al., 2013; Wientjes et al., 2013). Over a 4-year record of Chl *a/b* data, however, the values show some seasonal variations; in winter, the values at 2am are slightly higher (2.13-2.21) than at 15 and 26 m (2-2.06), while in summer are even lower (Chl *a/b* around 1.7-1.8) and stable at all depths **(Suplemental Figure 3A**). These data suggest that LHCII is present in large amounts in this plant at all depths.

**Figure 3:**
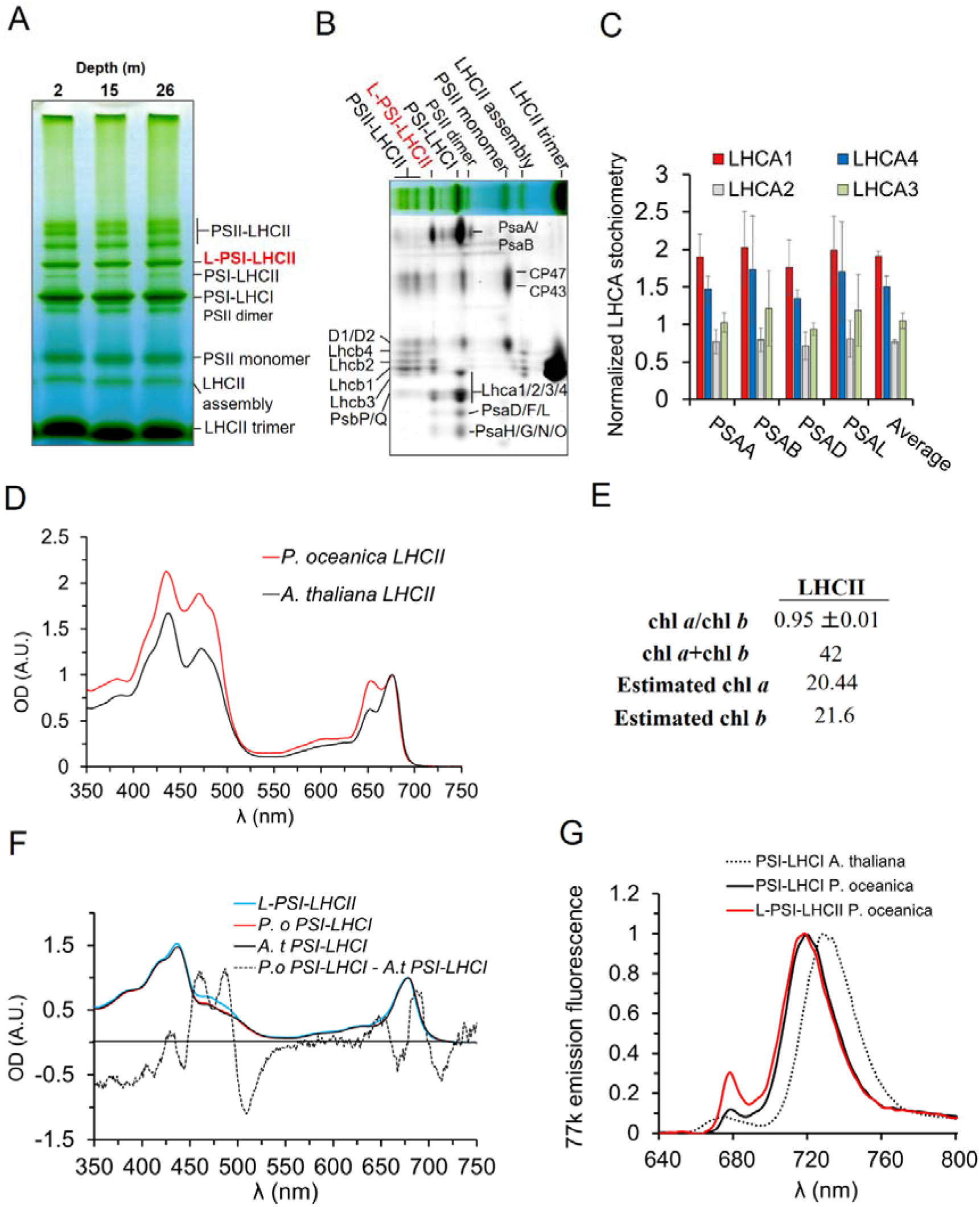
Analysis of thylakoid composition and the Large PSI-LHCII supercomplex. (**A**) BN-PAGE of solubilized thylakoid collected at different growing depths in winter. For each tested depth, 5μg of was loaded onto the gel. **(B)** Identification of the peptide content of the BN-PAGE bands by a second dimension on Urea-PAGE. Gel stained using SyproRuby **(C)** Proteomic analysis (LC-MSMS) of L-PSI-LHCII. Results were compared to PSI–LHCI. A number of peptide spectra matches (PSMs) are represented as fold change of the unique peptides from the 4 LHCAs normalized by the PSM of some of the PSI-core unique-peptides (indicated on the x-axis) between the L-PSI-LHCII and the PSI-LHCI. **(D)** Absorption spectrum of LHCII trimers purified from *P. oceanica* and *A. thaliana*. **(E)** Chl content of *P. oceanica* LHCII **(F)** Normalized (680 nm peak) absorption spectrum of PSI-LHCII compared to PSI-LHCI from *P. oceanica* and *A. thaliana*. The absorption difference spectrum between PSI from *P. oceanica* and *A. thaliana* was multiplied by 50 for better visualization. **(G)** 77K fluorescence emission spectra of PSI-LHCI and L-PSI-LHCII compared with the PSI-LHCI of *A. thaliana*, normalized at the 720 nm band maxima.

### The PSI/PSII ratio shows slight variation throughout depth

Since the difference in light spectra at different depths can result in different excitation of the two photosystems, we investigated if changes occur in PSI/PSII ratio with depths. We first assessed the PSI and PSII stoichiometry via immunodetection of PSII (CP43, CP47) and PSI (PsaG, Lhca2) subunits. The analysis of the blots (**Figure 2B and suplemental Figure 4A**) could not highlight significant variations, irrespective of the growing season of the collected sample (**Suplemental Figure 4B).** We then quantified the functional PSI/PSII stoichiometry using flash-induced electrochromic shift (ECS) at 520 nm on fresh untreated and DCMU infiltrated leaves (Bailleul et al., 2010). The stoichiometry variation was difficult to assess due to the deviation between experiments and replicates (**Suplemental Figure 4D and 4E)**, yet, at all depths tested, *P. oceanica* exhibits a PSI/PSII stoichiometry around 1.15 ± 0.06 (**Figure 2C**), higher than most land plants such as *A. thaliana* (0.6 to 0.9 grown under white light and sunlight, respectively (Hogewoning et al., 2012; Wientjes et al., 2017).

**Figure 4:**
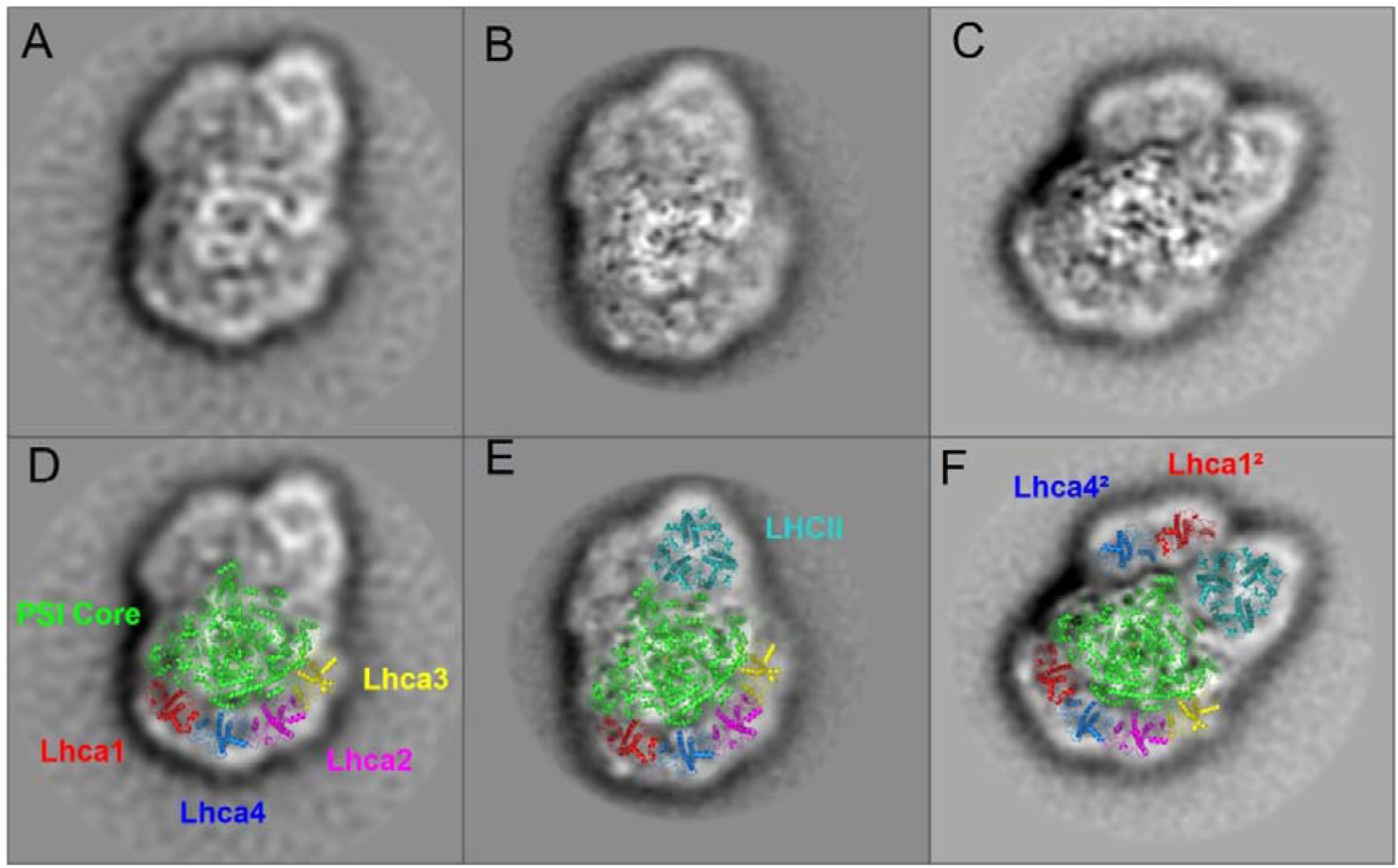
Architecture of the L-PSI-LHCII. **(A-F)** Negative staining single particles analysis of the L-PSI-LHCII purified from *P. oceanica*. **(a)** 2D EM projection class average size = 1441 particles (**b**) 2D class average size = 2787, (**c**) 2D class average size = 2629 particles. (**D-F**) The atomic models of the existing structure (5ZJI) were superposed on the L-PSI-LHCII projection 2D class averages: PSI core (green), Lhca1 (red), Lhca2 (magenta), Lhca3 (yellow), Lhca4 (blue) and LHCII trimer (cyan).

### Analysis of PSI and PSII supercomplexes and characterization of a Large PSI-LHCII supercomplex

Insights into the composition of PSI and PSII supercomplexes were obtained through the BN-PAGE analysis of *P. oceanica*’s α-DDM-solubilized thylakoids from different depths (**Figure 3A).** A distinct and intense band in between the PSII-LHCII bands and the PSI-LHCI band was observed. 2D urea-PAGE analysis shows that this band contains the main protein components of PSI-LHCI, with in addition LHCII subunits (Lhcb1 and Lhcb2) like the PSI-LHCII state transition complex in *A. thaliana* (Wu et al., 2023) and *Z. mays* (Pan et al., 2018). This Larger PSI-LHCII was designated as L-PSI-LHCII and its composition was analyzed by mass spectrometry.

The LC-MS-MS data showed that L-PSI-LHCII has a higher relative content of Lhca1 (around 1.8) and Lhca4 (around 1.5) peptides in comparison with PSI-LHCI (**Figure 3C),** indicating the presence of an additional Lhca1-Lhca4 dimer (Lhca1²-Lhca4²) bound to PSI-LHCII.

As for the composition of the LHCII trimer, we found a 2:1 stoichiometry of Lhcb1 and Lhcb2 (**Suplemental Figure 5 A-C**), similar to *A. thaliana* and maize, as well as threonine phosphorylation of the Lhcb2 subunit. L-PSI-LHCII and PSI-LHCI bands exhibit consistent intensity across all depths, regardless of the growing season (**Figure 3A**, **Suplemental Figure 6 and 7**). Quantification of the Native-PAGE bands reveals a L-PSI-LHCII/PSI-LHCI stoichiometry of 0.4, so the large complex represents at least 30% of the total PSI pool at all depths. L-PSI-LHCII seemed stable as a very weak amount of phosphorylated Lhcb2 protein was found in the free LHCII bands in the Native-PAGE (**Suplemental Figure 5D**). Moreover, phosphorylation levels of Lhcb2 in thylakoids show no significant variations between depths, consistent with the constant accumulation of the L-PSI-LHCII across depths when samples are collected during the day, regardless of the season of collection **(Suplemental Figure 5E)**. In contrast, plants harvested before sunrise have a lower concentration of this complex (**Suplemental Figure 8**).

**Figure 5:**
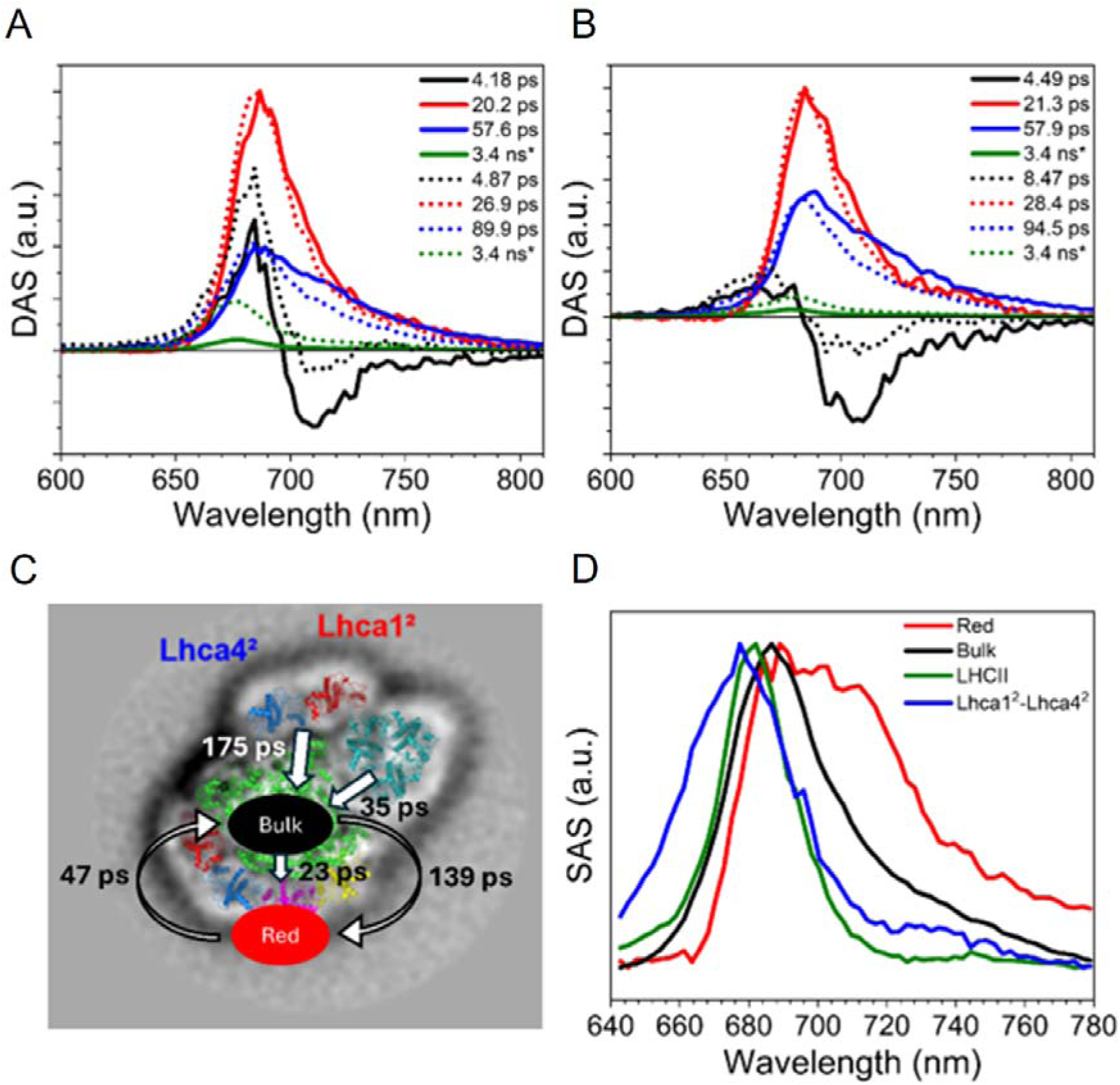
Excitation energy transfer and trapping of PSI-LHCI and L-PSI-LHCII. Decay-associated spectra (DAS) for the measurement of the PSI-LHCI sample (solid lines) and L-PSI-LHCII sample (short-dots) upon 400 nm **(A)** and 475 nm **(B)** excitation. The * indicates a fixed lifetime component. **(C)** Simplified kinetic model for the target analysis, see **Suplemental Figure 12** for the full scheme and an accompanying descriptive text. **(D)** Species-associated Spectra (SAS) as extracted from the target analysis. Bulk Chls are in black and the red forms in red.

**Figure 6:**
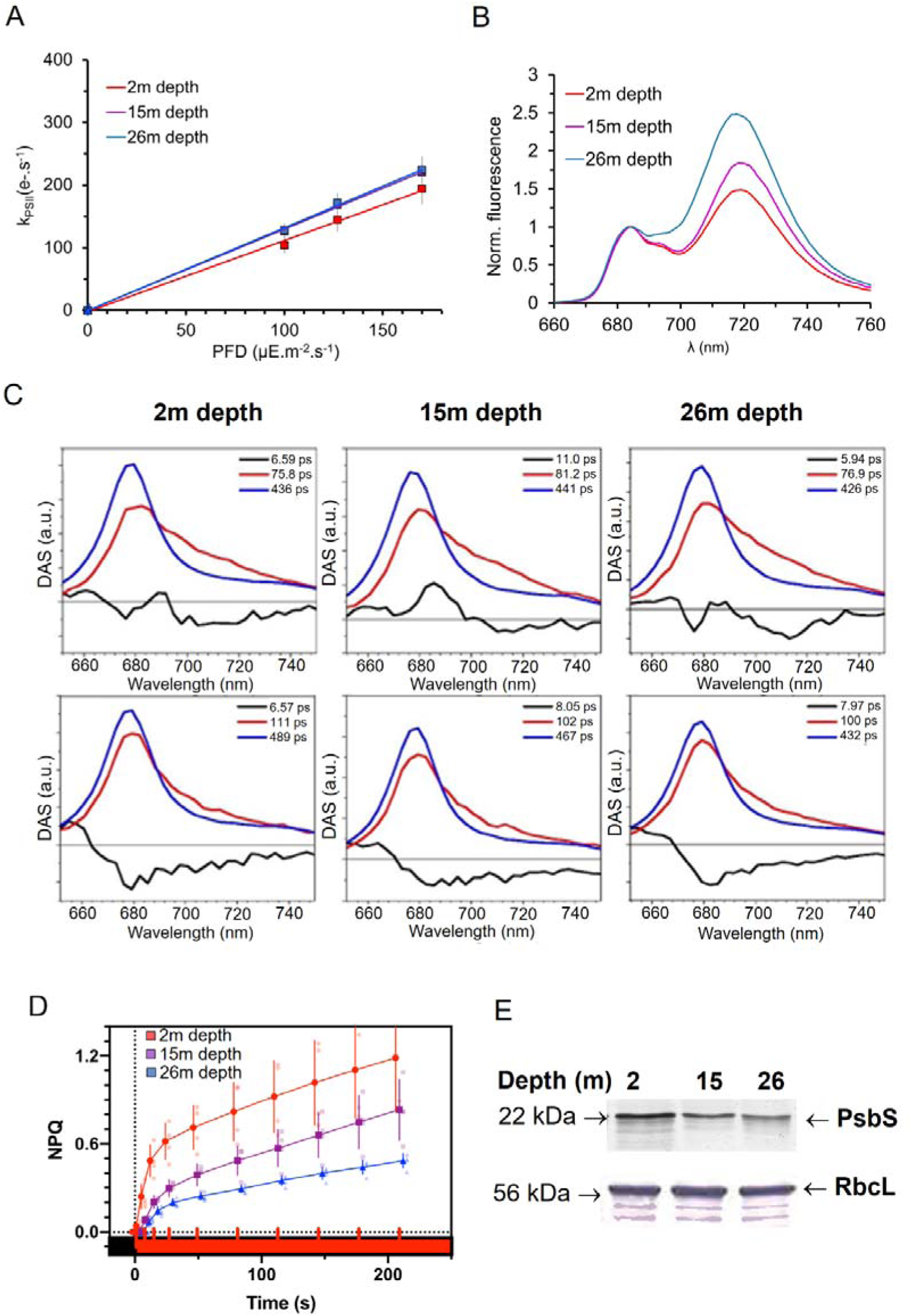
PSI and PSII light harvesting in *P. oceanica* at different growing depths. (**A**) Functional antenna size of PSII. PSII turnover rates (in e^−^.s^−1^), derived from the fluorescence rise kinetics of DCMU-blocked leaves collected in winter, were measured at three moderate (100, 125, and 170 µmol.m^−2^.s^−1^) intensities of red light. Slope values measure PSII functional antenna size. **(B)** Normalized 77K emission fluorescence of thylakoids from leaves collected at depths of 2, 15, and 26 m in winter**. (C)** Decay-associated spectra obtained from global fitting of time-resolved fluorescence spectra. Thylakoids from leaves collected at depths of 2, 15, and 26 m in winter were excited at 400 (top) and 475 nm (bottom). **(D)** Kinetics of NPQ induction. NPQ is probed on low light-adapted *P. oceanica* leaves exposed to moderate actinic illumination (200 µmol.m^−2^.s^−1^) n=3. **(E)** Immunodetection of PsbS in *P. oceanica* leaf exctract. Large rubisco subunit was used as a standard for protein loading.

**Figure 7:**
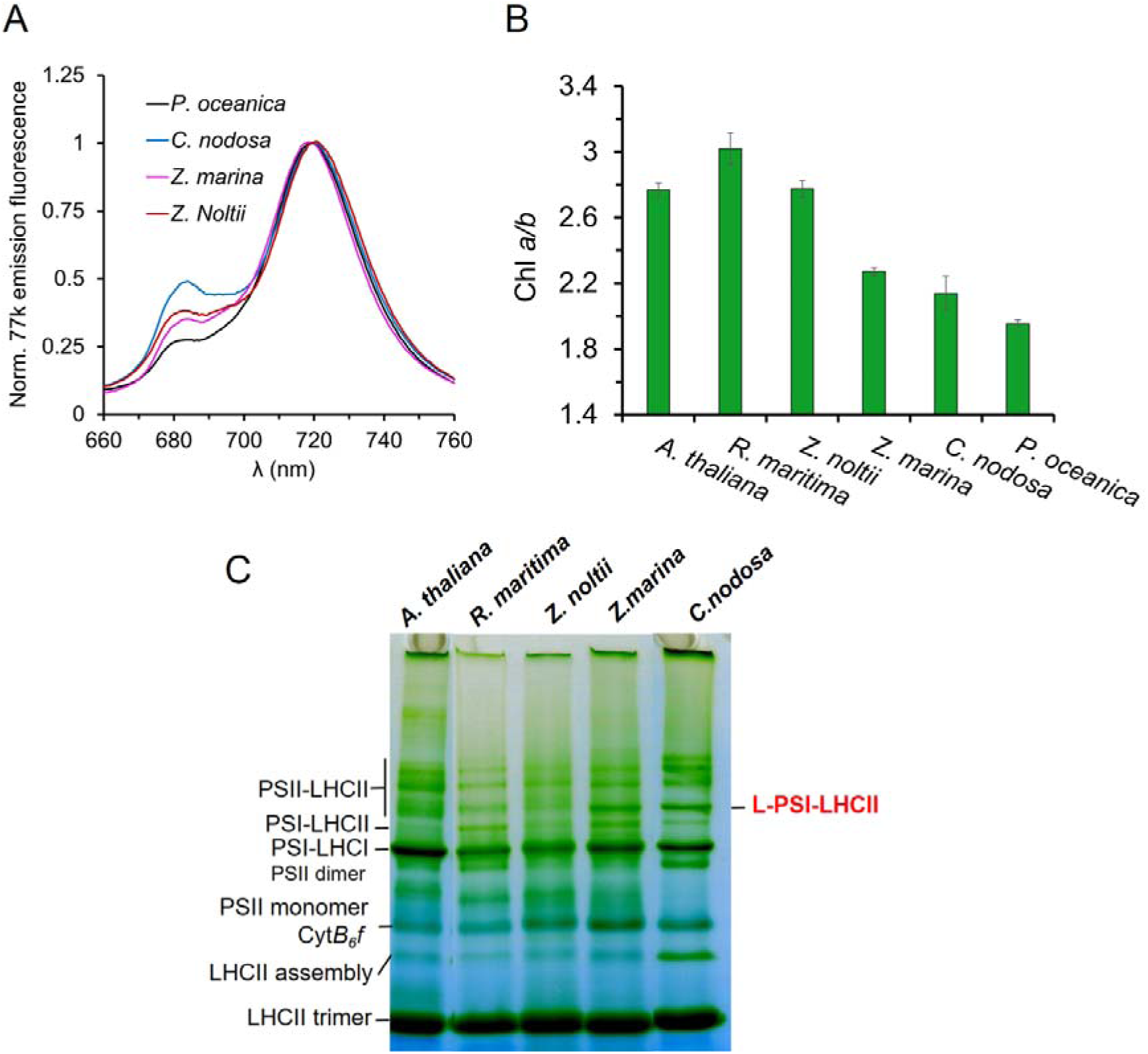
Thylakoid composition of different marine plant species. (**A**) 77K fluorescence emission spectra of the isolated PSI-LHCI from the selected plant species (indicated) normalized at 720 nm. The complexes were isolated by CN-PAGE. **(B)** Chl *a/b* of the thylakoid membranes from the selected plants. Data are expressed as the mean ± SD, n = 3. **(C)** BN-PAGE of solubilized thylakoids extracted in different seagrass species.

### Chlorophyll *b* enrichment in the LHCII, PSI-LHCI and L-PSI-LHCII complexes

Next we isolated the main pigment-protein complexes using sucrose density gradient ultracentrifugation. The absorption spectrum of *P. oceanica* trimeric LHCII showed higher amplitudes in the 450-500 nm and 650 nm regions compared to that of *A. thaliana* (**Figure 3D**), indicating a higher Chl *b* content. This was confirmed by pigment analysis, which showed a Chl *a/b* ratio of 0.95 土0.01, (**Figure 3E**). In the model organisms *Chlamydomonas reinhardtii*, *Physcomitrella patens, A. thaliana*, and *Zea mays*, the LHCII trimer contains 42 Chls with a Chl *a/b* ratio of 1.33 and therefore binds 24 Chl *a* and 18 Chl *b* (Huang et al., 2021; Zhang et al., 2023; Wu et al., 2023; Pan et al., 2018). Assuming the same total number of pigments, *P. oceanica* LHCII is expected to bind 20.5 Chl *a* and 21.5 Chl *b* (**Figure 3E**), suggesting that 3 Chl *a* have been replaced by Chl *b*.

The comparison of *P. oceanica* PSI-LHCI and the L-PSI-LHCII absorption spectra with that of *A. thaliana* PSI-LHCI (**Figure 3F**) shows a Chl *b* enrichment in the seagrass. This was confirmed by measuring the Chl *a/b* ratio of 6.84 士 0.05, and 4.33 ±0.03 in the PSI-LHCI and L-PSI-LHCII respectively, which are clearly lower than those of *A. thaliana* (**Suplemental Figure 9**).

In addition, the absorption in the far red of the PSIs from *P. oceanica* is less intense than in *A. thaliana* (**Figure 3F and Suplemental Figure 10**), indicating that these complexes contain less low energy pigments. This is confirmed by the 77K emission fluorescence spectra that for both PSI-LHCI and L-PSI-LHCII show a peak at 718-719 nm, significantly blue-shifted compared to PSI-LHCI of *A. thaliana* (735 nm) (**Figure 3G**).

### Structural organization of L-PSI-LHCII

To gain further insight into the structural organization of the L-PSI-LHCII, we performed single-particle analysis using negative staining electron microscopy (**Figure 4**). The 2D class average projections show a density corresponding to PSI-LHCI, one to trimeric LHCII and an extra density corresponding to the additional Lhca1-Lhca4 dimer. The Lhca1-Lhca4 dimer interacts PSI-core on the PsaB side (**Figure 4D-F**), while the LHCII trimer is connected to the PsaA, side, as observed in land plants (Wu et al., 2023).

### Energy transfer dynamics in the PSI-LHCI and L-PSI-LHCII complexes

To investigate excitation energy transfer (EET) and trapping in *P. oceanica* PSI-LHCI and L-PSI-LHCII complexes, we performed time-resolved fluorescence measurements with 400 nm and 475 nm excitation, which excite the core and the outer antennae differently (see **Suplemental Figure 11 for** streak camera images). Global analyses were performed on these datasets and four components were sufficient to fit each dataset satisfactorily (**Figure 5A-B**). The fast component (<10 ps) represents EET from high to low-energy Chls, while the ns component, with small amplitude and a blue-shifted spectrum corresponds to disconnected Chls. The dominant ∼20 ps (in PSI-LHCI) and ∼28 ps (in L-PSI-LHCII) component, has a similar spectrum in all datasets and is associated with trapping excitations in the RC from bulk Chls, mainly in the PSI core. A red-shifted 58 ps (in PSI-LHCI) and 94 ps (in L-PSI-LHCII) component, with larger amplitude upon 475 nm excitation, describes trapping in the RC from excitations mainly originating in the antenna. The longer lifetime in L-PSI-LHCII reflects its larger antenna. Notably, the second component also shows a slight increase in lifetime in L-PSI-LHCII (from ∼20 ps to ∼28 ps) and relative amplitude, suggesting the presence of an antenna complex without red forms that transfers energy rapidly to the PSI core.

To better understand the EET dynamics, we simultaneously fitted the four datasets using a target model (summarized in **Figure 5C**). The full kinetic scheme and fitting quality are presented in **Suplemental Figure 12**. Target analysis reveals that excitations in the bulk Chls are trapped in 23 ps, consistently with previous studies on different organisms (Engelmann et al., 2006; Wientjes et al., 2011; Le Quiniou et al., 2015) as expected as the PSI core is highly conserved among plants, algae and cyanobacteria (Alboresi et al., 2017). EET from red forms to bulk Chls occurs in 47 ps, while the reverse transfer has a lifetime of 139 ps. This is a consequence of the significantly higher amount of bulk Chls than red forms in the system, providing an entropic advantage to this compartment, although it is higher in energy. The Species-Associated Spectra (SAS) (**Figure 5D)** confirm the attribution of the compartments to bulk Chls and red forms. The extra LHCII trimer is energetically reasonably well connected to the bulk Chls (forward time constant of 35 ps and a reverse time constant of 686 ps). The extra Lhca14 dimer is instead less well connected to the bulk: it exchanges energy with the bulk at a forward rate of (175 ps)^−1^ and a reverse rate of (819 ps)^−1^.

### The antenna size of both Photosystems remains constant with depth

The low Chl *a/b* ratio of *P. oceanica* suggests that more LHCII than in e.g *A.thaliana* are present in the membrane. Using the method from (Ünlü et al., 2014) we could estimate that there should be between 6 and 8 LHCII trimers per PSII, with only minor changes between depths (**Suplemental Figure 3B**).

The functional antenna size of PSII was determined following the fluorescence rise kinetics in DCMU-treated leaves (Erickson et al., 1984) (**Figure 6A**). The Antenna size was slightly lower at 2 m compared to 15/26 m but, overall, it appeared highly similar at all depths (**Figure 6A**), indicating that a change in PSII antenna size is not a major adaptation strategy to depth in *P. oceanica*. Next, we tested the the distribution of LHCII between the two photosystems. The 77K fluorescence emission spectra of the isolated thylakoids showed a depth-dependent increase of the 715 nm emission relative to 680 nm (**Figure 6B**). These differences can have various origins as the signal depends not only on the number of pigments associated with the two photosystems but also on their lifetimes (in this respect is important to consider that PSI lifetimes becomes long and multiexponential at 77K) and the connections between the complexes. Thus, both the composition and organization of the thylakoids could explain the changes in fluorescence emission with depth (Rantala et al., 2017).

For a more quantitative approach, we performed time-resolved fluorescence measurements on winter (**Figure 6C**) and summer thylakoids (**Suplemental Figure 13A**). This technique allows us to disentangle the contributions of the two photosystems. The lifetime of PSI will inform about the antenna size of the complex and the area under the components of PSI and PSII proximates the relative number of pigments associated with the two photosystems. Exiting at 400 nm and 475 nm provide information on the migration time from the antenna to the PSI core.

Three components were sufficient to fit the datasets (**Figure 6C and Suplemental Figure 13A**). The first component (<10 ps, black in **Figure 6C**) represents EET from blue to red chlorophylls in PSI corresponding to the fast component in the isolated PSI complexes (**Figure 6C**). The second component (red in **Figure 6C**) exhibits the far-red signature of PSI and has similar lifetimes and spectra (**Suplemental Figure 13B** for PSI DAS comparison) across depths, indicating no differences in PSI antenna size. However, both spectra and lifetimes are excitation wavelength-dependent. The difference in PSI lifetime between excitations (∼100 ps at 475 nm and ∼80 ps at 400 nm) indicates that the LHCs (which contain Chl *b*) significantly contribute to the PSI antenna with some complexes transferring energy relatively slowly to the RC. This agrees with the measurements on the isolated L-PSI-LHCII complex. Moreover, the PSI spectrum upon 475 nm excitation shows a relatively larger amplitude at 680 nm than above 700 nm, (**Suplemental Figure 13**), indicating that the outer antenna complexes mainly contribute to the emission at 680 nm, while the far-red emission (>700 nm) originates from the core. The third component (blue in figure 6) also has similar lifetimes and spectra in all datasets and is associated with PSII. While lifetimes and spectra remain similar at different depths for all components, depth-dependent changes are observed in the relative areas of the components associated with PSI and PSII, increasing slightly with depth with the largest variation (from 0.9 to 1.1) in summer leaves. This result indicates a change in the excitation between PSI and PSII, resulting from a difference in the number of Chls associated with the two photosystems at different depths. Considering that the PSII antenna size does not change (**Figure 6A**), there are still two remaining explanations for this result: (i) a change in the PSI antenna size and (ii) a change in PSI/PSII ratio. Since the PSI lifetime and spectra remain unchanged with depths, indicating no changes in PSI antenna size, we conclude that these differences are due to a small increase in the PSI/PSII ratio from 2 m to 26 m depths.

### Non-photochemical quenching and Psbs expression at different growth depths

Balancing the energy distribution between photosystems may also occur through modulation of their photochemical efficiency (ultimately affecting their functional antenna size). Shallow *P. oceanica* leaves showed a ∼3 times higher non-photochemical quenching (NPQ) than 26 m depth leaves, while 15 m depth leaves displayed an intermediate NPQ level (**Figure 6D**). PsbS expression followed this pattern (**Figure 6E**), showing an increase in shallow/high-light plants, and conversely a strong decrease in deep/low-light plants. Like *A. thaliana*, PsbS expression is light-depending (Albanese et al., 2016) and NPQ depends on the amount of the PsbS protein (Li et al., 2002).

### L-PSI-LHCII complex in other Mediterranean seagrasses

The presence of the L-PSI-LHCII was investigated across four mediterranean seagrass species with different depth growth limits: *Cymodocea nodosa* (1-40 m), *Zostera marina* (up to 15 m) and *Ruppia maritima*, and *Zostera noltii* (up to 2 m). All species were harvested at 0.5-1 m depth. 77k fluorescence spectra of isolated PSI revealed a maximum emission peak at 720 nm for all species (**Figure 7A**). Comparing Chl *a/b* ratios in purified thylakoids showed that the shallow plants *R. maritima* (3.01) and *Z. noltii* (2.77) have a ratio higher than *P. oceanica* grown at similar irradiance, whereas *C. nodosa* (2.14) is closer to *P. oceanica* (1.95), suggesting a higher LHCII abundance in species with wider depth distribution. Z*. marina* (0.5-15 m) had an intermediate Chl *a/b* ratio (2.27) between shallow and deep seagrasses (**Figure 7B**).

BN-PAGE was performed to analyze thylakoid complexes in the selected plant species. *Z. marina* and *C. nodosa* showed a strong green band, matching the molecular weight of L-PSI-LHCII (**Figure 7C**). In contrast, *Z. noltii* and *R. maritima* exhibited migration profiles similar to *A. thaliana*, as the position of the L-PSI-LHCII is occupied by PSII supercomplexes as seen by the 2D-PAGE analysis (**Suplemental Figure 14**) suggesting the absence of L-PSI-LHCII in plants specifically thriving in shallow waters. To confirm the absence of L-PSI-LHCII in *A. thaliana* and shallow seagrasses, the Lhca4 protein signal was compared to that of Lhca2 in *P. oceanica*, *A. thaliana*, *R. maritima*, and *Z. noltii* using immunoblotting (**Suplemental Figure 15**). The Lhca4/Lhca2 ratio from the blots was normalized to the Lhca4/Lhca2 ratio in the purified PSI-LHCI, where the stoichiometry between both proteins was assumed to be 1. In *P. oceanica*, the ratio was 1.43 ± 0.06, while in *R. maritima*, *Z. noltii*, and *A. thaliana*, it was 1.02 ± 0.06, 0.99 ± 0.07, and 1.07 ± 0.07, respectively. This indicates that seagrasses with a shallow distribution, similar to the land angiosperm *A. thaliana*, have only a few/no PSI complexes with an additional Lhca1-Lhca4 antenna. In contrast, the strong accumulation of L-PSI-LHCII is a characteristic of seagrasses able to colonize a wider bathymetric area.

## Discussion

Chl *b* within LHCII, primarily associated with PSII, augments its absorption in the blue-green spectral region, whereas red and far-red photons preferentially excite PSI. In alignment with the blue-enriched spectral environment, *P. oceanica* displays a PSI/PSII stoichiometry of around 30-40% relative to *A. thaliana*, compensating for PSI’s inherent absorption deficit. Beyond the overall PSI content, *P. oceanica* exhibits a high LHCII expression compared to *A. thaliana*. Furthermore, the PSI-LHCI complex and the LHCII exhibit an increased Chl *b* enrichment, thereby enhancing the absorption of blue wavelengths. The augmentation of PSI is manifested through the formation of L-PSI-LHCII supercomplexes, representing 30% of the total PSI pool, and which is characterized by the incorporation of a chlorophyll b-enriched LHCII trimer (Figure 3) alongside an additional Lhca1-Lhca4 heterodimer.

In land plants, red forms located in Lhca3 and Lhca4 absorb light below the P700 reaction center energy. Although they are essential for harvesting far-red light, they slow PSI’s energy trapping (Engelmann et al., 2006; Jennings et al., 2013; Chukhutsina et al., 2020). In marine environments depleted of far-red light, *P. oceanica* lost those red-forms, allowing it to increase its PSI antenna size, through the connection of the extra Lhca1-Lhca4 dimer, while preserving high energy-trapping efficiency (**Figure 5**). Additionally, the longer trapping time of *P. oceanica*’s PSI in the membrane compared to L-PSI-LHCII (100 ps vs. 55 ps at 475 nm excitation) suggests that PSI *in vivo* includes at least one more LHCII trimer, possibly lost during purification as seen in other studies (Croce, 2020) (**Figure 6, Suplemental Figure 12 and 13**). It can be hypothesized that the L-PSI-LHCII could serve as an anchor point for the binding of one or multiple LHCII, notably via the canonical and/or extra Lhca4 subunits as suggested in *A.thaliana* (Benson et al., 2015), with the absence of red forms in *P. oceanica* which could maybe facilitate energy transfer efficiency from the extra LHCII to the core.

The L-PSI-LHCII supercomplex has been consistently identified in *P. oceanica* and three additional marine alismatales species (*C. nodosa*, *Z; marina*, and *H. stipulacea*), all of which exhibit extensive bathymetric distributions (**Figure 7, Supplemental Figure 16**). In contrast, this complex was not detected in seagrass species adapted to shallow photic zones. A comparable assembly has been reported in *A. thaliana* (Crepin et al., 2020); however, it lacked Chll *b* enrichment, did not exhibit the characteristic absence of red forms, and appeared to be present at levels too low to fulfill a substantial light-harvesting function. Furthermore, we showed that a decreased chlorophyll *a/b* ratio in the thylakoid membranes of the tested seagrasses correlates with their bathymetric distribution and the presence of the L-PSI-LHCII. Collectively, these data strongly indicate that the L-PSI-LHCII supercomplex enhances low-blue light-use efficiency by: (i) increasing the antenna cross-section via an additional Lhca1-Lhca4 dimer alongside LHCII, (ii) enriching chlorophyll b to boost blue light harvesting, (iii) lacking red-shifted forms which enables faster excitation energy trapping in PSI compared to land plants, and (iv) potentially binding additional LHCII units. Furthermore, similar strategies such as a high PSI/PSII ratio, high LHCII content together with an enrichment of Chl *b* in the LHCII, are also expressed in green marine macroalgae *Bryopsis maxima* and *Ulva pertusa* (Yamazaki et al., 2005), highlighting an example of convergent evolution between green algae and seagrasses.

Surprisingly, L-PSI-LHCII and, to a larger extent, both PS’s antenna size, as well the PSI/PSII ratio in *P. oceanica* remain rather stable at all depth. In contrast, as seen in other organisms (Zuo, 2025), we showed a greater NPQ activation in shallow plants, with this capacity varying by depth through PsbS protein expression level (**Figure 7**). Additionally, microscopy observations showed that thylakoid’s organisation changed drastically (**Figure 1**) as a consequence of the depth-specific light constraints. Specifically, the reduced grana size observed at 2 meters depth may enhance PSII repair by facilitating the mobility of light-damaged PSII complexes toward the grana margins to facilitate repair mechanisms as suggested by (Procaccini et al., 2017). Conversely, at 26 meters, a wider yet thinner grana combined may facilitate light penetration into the grana’s internal stacks. A comparable structure has been already observed in *A. thaliana* where it would allow greater efficiency for linear electron transfer under low light (Garty et al., 2024). Nevertheless, (Procaccini et al., 2017) showed that the linear electron transfer (LET) measured *in situ* in *P. oceanica* decreases in deeper meadows as a consequence of reduced irradiance. All together, these observations suggest that photosynthetic efficiency in *P. oceanica* is primarily influenced by light intensity and quality, as well as the organization of thylakoid membranes, rather than solely by antenna size or photosystem stoichiometry.

We propose that *P. oceanica* maintains its photosynthetic apparatus in a configuration maximizing light harvesting at all depth. Yet, a dynamic modulation of antenna connectivity notably through NPQ, particularly in shallow waters, would be essential for coping with fluctuating light intensities throughout the day (Procaccini et al., 2017). A complementary explanation is that as depth increases, canopy density decreases, partially compensating for the reduced light availability. Conversely, in shallower meadows, leaf density results in self-shading (Dalla Via et al., 1998) and they are more prone to colonization by epiphytic organisms (Tsirika et al., 2007), meaning that the actual light received by the leaves is lower than the incident irradiance measured (**Suplemental Figure 1**). The reduced variability in *P. oceanica* may also reflects the depth-dependant need to sustain energy production. In shallower waters, plants are exposed to higher light intensity but also face increased energy requirements, related to *e.g* structure or defense, while in deeper waters, lower light availability may be balanced by reduced energy needs (Olesen et al., 2002; Dattolo et al., 2013; Costa et al., 2015; Ismael et al., 2023; Charras et al., 2024). In all cases, these energetic needs likely remain higher than those typically experienced by plants under controlled laboratory conditions. Hence, regulatory patterns observed in lab-grown species may not fully reflect the photosynthetic strategies express in the wild to cope with complex and fluctuating environments, whether marine or terrestrial.

The photosynthetic organization observed in *P. oceanica* appears to reflect a finely tuned balance, where energy demands across depths have selected for a stable and low-plasticity photosynthesis. While this constancy may be successful in stable marine environments, it could now become a vulnerability under rapid anthropogenic change. Since the mid-20th century, coastal urbanization has degraded *P. oceanica* habitats and increased water turbidity, limiting its bathymetric range to ∼30Lm near urbanized areas. Rising sea levels are expected to further reduce light availability at the lower limit of meadows. In parallel, the invasive species *Halophila stipulacea* is spreading rapidly. Even though it expresses the L-PSI-LHCII, unlike *P. oceanica*, *H. stipulacea* does not form a persistent matte crucial for long-term carbon storage. Effective conservation must therefore focus on protecting existing meadows, improving water transparency to support deeper recolonization, and controlling *H. stipulacea*’s spread. Collectively, these efforts will help sustain and possibly expand this essential marine carbon sink.

## METHODS

### Sample collection

*P. oceanica* shoots were collected from three healthy meadow patches at 2, 15, and 26Lm depth near Frioul Island, Marseille, France (43.1611LN, 5.1732LE), avoiding the fragile upper (0.5Lm) and lower (29Lm) limits. Sampling occurred in January and June of 2019– 2022 between 9–11Lam. Plants were kept at 4 °C in the dark and processed in the lab within 3 hours: leaves were separated at 3 °C under green light, and epiphytes removed by seawater washing. *C. nodosa* and shallow *P. oceanica* were collected in Agay (43.4299LN, 6.8680LE) in June 2021 (0.5Lm depth), and *Z. marina* and *Z. noltii* in Port de Bouc (41.2158LN, 4.5152LE) in July 2021 (0.5Lm depth). A. thaliana WT Col-0 plants were grown for 4 weeks under 120 µmol photonsLm⁻²Ls⁻¹ white light.

### Isolation and separation of photosystem supercomplexes

Thylakoid membranes were prepared as described in (Charras et al., 2024).

### Thylakoids membrane preparation for separation of photosystem supercomplexes by Blue-Native-PAGE

Thylakoid membranes were washed 2 times in a solubilization buffer (50 mM BisTris pH=7, 25 mM EDTA, 20% glycerol, 10 mM NaF). The chlorophyll *a/b* ratio and chlorophyll concentration were determined according (Croce et al., 2002; Chazaux et al., 2022). At 4°C, purified thylakoids (1 mg/ml) were solubilized with a final concentration of 2% (v/w) n-Dodecyl-L-D-Maltopyranoside solution (α-DDM, Anatrace, Ref: D310HA 10 GM) for 20 min, then centrifuged at 21,000 g. The solubilizate was then used for Blue native-PAGE sample preparation.

### Separation of photosystem supercomplexes by Blue native-PAGE

As in Järvi et al., 2011^37^, the solubilizate was supplemented with 1/10 volume of Serva Blue loading buffer (100 mM BisTris pH 7.0, 0.5 M amino-caproic acid, 30% (w/v) sucrose and 50 mg/mL Serva Blue G). About 5-10 μg of chlorophyll was then loaded on the commercial gel (Invitrogen™ NativePAGE™ 4 –16L%, Bis-Tris, 1.0 mm; ref BN1001BOX) set in Minicellule XCell *SureLock*™ Invitrogen™. The anode buffer contained 25 mM BisTris, 25 mM tricine pH=7, and the cathode buffer contained 50 mM BisTris pH=7, 0.015% LDDM, and 0.01% Serva Blue G250. Electrophoresis was performed at 4°C for 7 hours. The intensity was fixed and set so that at the start of the run, the voltage should be around 40V.

### Second Dimension urea-PAGE identification of polypeptides content from supercomplexes isolated by BN-PAGE

The lanes of interest were cut from the 1D BN-PAGE and incubated for 30 min in Laemmli denaturing buffer (125 mM Tris pH 6.8, 20% glycerol, 2% SDS, 100 mM DTT, 0.01% bromophenol blue). The lane was then deposited on the top of 13% acrylamide/bisacrylamide (37.5/1) resolving gel containing 6 M urea and fixed with agarose (1% agarose, 0. M Tris pH 6.8, 0.001% bromophenol blue). The anode and cathode chambers were filled with a Laemmli running buffer (25 mM Tris pH 8.3, 0.192 M glycine, 0.1% SDS). Migration was performed for approximately 2h30 at 90 V.

### Urea-PAGE and immunoblot

Thylakoid preparations were denatured at 70 °C for 30 min in denaturing Laemmli buffer, then loaded on a 14% acrylamide gel (37.5:1 acrylamide/bisacrylamide) containing 6LM urea (Tris-Glycine Laemmli system). For Phos-tag™ analysis, 12% gels with 50 μM Phos-Tag (Wako Pure Chemicals Industries) and 100 μM MnClL were used. After electrophoresis, gels were stained with Sypro™ Ruby or used for immunoblotting. Phos-tag™ gels were equilibrated in transfer buffer with 50 mM EDTA before transfer using the BioRad Trans-Blot Turbo system. Immunoblots were developed using Agrisera antibodies and chemiluminescence detection (Fusion FX7, Vilber). Leaf extracts were prepared as described in (Charras et al., 2024).

### L-PSI-LHCII and LHCII purification

Purified thylakoids (1 mg/ml) were solubilized (same buffer as Native-PAGE without glycerol) with a final concentration of 2% (v/w) U-DDM for 20 min, then centrifuged at 21,000 g. The supernatant was loaded onto a sucrose gradient (0.75 M sucrose, 40 mM Hepes pH 7.4, 0.015% U-DDM), formed by freeze-thawing. After centrifugation, target bands were collected, dialyzed against 40 mM Hepes pH 7.4 concentrated. For LHCII, the sample underwent a second sucrose gradient, followed by dialysis and concentration. For L-PSI-LHCII, PSI-LHCI contaminants were partially removed after the first gradient using anion exchange chromatography (MonoQ, 0–500 mM NaCl in 40 mM Hepes pH 7.4, 0.015% U-DDM).

### Electron microscopy analysis of chloroplast ultrastructure

Samples were prepared as in (Nja et al., 2018). Grids were observed under the Tecnai G2 electron microscope.

### Sample preparation for negative staining single particle electron microscopy analysis

5 μL of the protein sample (3 mg.mL^−1^) were applied to the glow-discharged (PELCO easyGlow, 15mA, 25 s) Quantifoil 100 Carbon Support Films grid: Cu 300 and left on the grid for 45 sec. The excess sample was blotted away (Whatman #1). 5 μl of 2% uranyl acetate was applied to the grid. Excess stain excess was blotted away (Whatman #1) for 30s. Grids were washed by applying a 5% uranyl acetate droplet (5 μl), removed immediately with blotting paper (Whatman #1). After drying out, the grids were loaded in the Thermo Fisher Scientific Glacios using an autoloader under cryo-EM conditions.

### Electron microscopy data acquisition

Data acquisition was performed on the Thermo Fisher Scientific Glacios equipped with a field emission gun and operated at 200 kV in bright field imaging mode. Movies were recorded using a Thermo Scientific Falcon 3EC Direct electron Detector in linear mode at a nominal magnification of 92,000, corresponding to a pixel size of 1.567 Å/pix with 30 frames at a dose of 30 e-/Å2 per frame and an exposure time of 30 s per movie.

### Image processing

The dataset of PSI protein (fraction p1q_15) was derived from 1326 movies. The raw movies were imported into SCIPION 3.0 (de la Rosa-Trevín et al., 2013) for the following processing steps. The movies were motion-corrected using the MotionCor2-1.4.7 protocol (Zheng et al., 2017) and the CTF estimation was done by the Gctf – 1.06 protocol (Zhang, 2016).

A total of 337227 particles were extracted after xmipp manual/auto particle picking^55^. After multiple rounds of 2D classification using cryoSPARC v3.0.1 protocol (Punjani et al., 2017), the main protein complexes of interest: PSI and PSI-LHCII, with apparent features, were sorted. Based on the 2D classification, the particles exhibiting the clear signatures of PSI-LHCI and L-PSI-LHCII were manually sorted into two groups: 1) PSI-LHCI, containing 168432 particles and 2) L-PSI-LHCII with 14255 particles.

### 77K fluorescence spectra

For the L-PSI–LHCII and PSI-LHCI from *P. oceanica* the supercomplexes were isolated as described above (sucrose gradient and AEC) and for the other plant species, the supercomplexes were isolated by CN-PAGE. The band was sliced and deposited in an aluminum holder and snapped frozen in liquid nitrogen at 77 K. By means of an optic fiber, chlorophyll is excited by a blue (∼455 nm) LED, and the resulting fluorescence signal is detected through a red-colored Kodak Wratten filter. Excitation and detection are mediated with an Ocean Optics USB2000 CCB spectrometer, piloted with an in-lab-developed PC acquisition software.

### Determination of PSI/PSII stoichiometry via ECS spectroscopy

In the green lineage, absorbance changes at 515–520 nm directly reflect thylakoid trasmembrane potential (Joliot and Joliot, 1989). A Joliot-type spectrophotometer (Continuum Nd:YAG Laser + OPO) was used with laser flashes (680 nm) for excitation, and BG39 filters to block actinic light. In *P. oceanica* leaves, a saturating flash induces a ΔA520Lnm peak, indicating membrane potential from one-electron transfer per photosystem (PSII + PSI). When PSII is inhibited (DCMU 10 µM + hydroxylamine 10 mM), a reduced peak reveals PSI’s sole contribution. The PSI/PSII ratio is then calculated from peak amplitudes (Bailleul et al., 2010). This method reflects only effective charge separation, independent of antenna size variations.

### Determination of PSII antenna sizes via DCMU-fluorescence rise

PSII functional antenna sizes have been determined using a Dual-PAM 100 fluorimeter (Melis and Homann, 1975; Lavergne and Trissl, 1995). In brief, the fluorescence rise (FR) kinetics was measured on *P. oceanica* leaves upon pulses (500 ms) of red light. Leaves were pre-treated with DCMU (to block all electron transfers downstream PSII) and kept in darkness at room temperature in fresh seawater. The area above the sigmoid curve (normalized from 0 to 1) is proportional to the still-oxidized primary acceptor Q_A_ fraction. The evolution of the Q_A_ reduction rate can be obtained by integrating the FR curve: the initial rate at light onset (slope at origin) gives the PSII functional antenna size.

### Non-photochemical quenching of fluorescence

Fluorescence kinetics were followed on fresh, low light-adapted *P. oceanica* leaves with a DualPAM-100 fluorometer. The maximal fluorescence yield *Fm* is first probed by a preliminary saturating light pulse on a dark background. After light onset, successive saturating pulses probe the evolution of the maximal fluorescence yield during photochemistry *Fm’*. The NPQ parameter is computed as (*Fm*-*Fm’*)/*Fm’* for each measurement.

### Time-resolved fluorescence spectroscopy

Time-resolved fluorescence measurements on *P. oceanica* samples were performed at room temperature using a synchroscan streak camera setup as previously described in Hu et al. (2023, New Phytol.). Femtosecond pulses at either 400 nm or 475 nm with a repetition rate of 250 kHz were used. The excitation spot size was ∼150 µm. An excitation power study was performed for each sample. The thylakoids were treated with 20 µM DCMU to close the reaction centers. The samples were magnetically stirred in 1 cm × 1 cm cuvette, and the OD at Q_y_ maximum was ∼0.5 for the thylakoids and ∼0.3 cm^−1^ for the PSI samples. The fluorescence was detected at a right angle using a spectrograph (Chromex 250IS, 50 l/mm) and a synchroscan streak camera (Hamamatsu C5680). Two-dimensional streak images were corrected for the detector’s background and wavelength-dependent sensitivity. The resulting averaged streak images were sliced up into time traces spanning ∼3 nm. Processed streak images were then globally analyzed using the Glotaran (Mullen and Stokkum, 2007; Snellenburg et al., 2012) and TIMP package for R to determine the fluorescence lifetimes and decay-associated spectra (DAS). Briefly, the streak images were fitted with a sum of four exponentials. The amplitudes of these exponentials across emission wavelengths resulted in decay-associated spectra.

### Target analysis

The streak images of the PSI-LHCI and L-PSI-LHCII samples excited at both 400 and 475 nm were simultaneously analysed in a target analysis using the pyglotaran Python package (Weißenborn et al., 2021; van Stokkum et al., 2023). The full kinetic scheme is presented in **Suplemental Figure 12**. To allow for the estimation of the initial excitation densities for the different compartments the areas under the SAS were constrained to be equal (Maiuri et al., 2013). The initial excitation densities are placed in the precursor compartments (precursor compartment names start with ET and they are grouped in the grey bar in **Suplemental Figure 12**. They were free fitting parameters for each measurement. The precursor compartments transfer rapidly (2.0-3.3 ps) and within the instrument response time (modeled as a Gaussian function with FWHM ranging from 5.3-6.7 ps) to the principal compartments, which are the ‘Bulk’, ‘Red’, ‘LHCII’ and ‘xLhca14’. The precursor compartments represent the fast energy equilibration within the subcomplexes and can be compared to the first (black) DAS of the global analyses. For each dataset/measurement the SAS of the precursor compartments were constrained to be equal. For each dataset/measurement there is also the ‘Disc.’ compartment to model the disconnected Chls. The measurements conducted on the PSI-LHCI samples were fitted with a model that for the principal compartments only contained the ‘Bulk’ and the ‘Red’ compartments, whereas for the measurements on the L-PSI-LHCII sample the model additionally contained the ‘LHCII’ and the extra Lhca1-Lhca4 dimer (Lhca1²-Lhca4²) compartment. The SAS of the principal compartments were constrained to be equal across all datasets. The backsweep of the synchroscan streak camera was also modeled and its frequency was fixed to 76 MHz. The fit quality of the target model is excellent and shown for a range of wavelengths in **Suplemental Figure 17**.

### Absorption spectrum deconvolution

For the spectral deconvolution the measured absorption spectra of the *P. oceanica* LHCII trimer, PSI-LHCI and L-PSI-LHCII complexes were first normalised to their integrated absorption in the Q_y_ region (630-750 nm) and then scaled to their relative Chl content as presented in **Suplemental Figure 9A**, taking into account the oscillator strength ratio of 1:0.7 for Chl *a*:*b*. The extra Lhca14 spectrum was obtained by subtracting the LHCII trimer and the PSI-LHCI absorption spectrum from the L-PSI-LHCII absorption spectrum.

### Mass spectrometry

Native-PAGE bands from the L-PSI-LHCII and *p.o* PSI-LHCI complexes were analyzed in triplicate by LC-MS/MS (Q-Exactive Plus, ThermoFisher), following (Proietti et al., 2023) with minor modifications. Samples were digested with Trypsin/Lys-C (Promega), and data processed using Proteome Discoverer 2.4.1.15 with the *A. thaliana* database (SwissProt, TxID 3702, version 2017-10-25) supplemented with *P. oceanica* reaction center and Lhca sequences. Peptides were validated via Sequest, using best PSM score (ΔCn ≤ 0.05) and 1% FDR. Spectral counting of unique peptide PSMs for Lhca proteins, normalized to core subunit PSMs, allowed estimation of relative Lhca abundance and L-PSI-LHCII/PSI-LHCI ratio.

### Declaration of generative AI and AI-assisted technologies in the writing process

During the preparation of this work the author(s) used ChatGPT in order to improve language and readability. After using this tool/service, the author(s) reviewed and edited the content as needed and take(s) full responsibility for the content of the publication.

## Author contribution

CJ initiated and developed the research project. CJ, and QCF planned and designed the experiments.

QCF performed sample preparation, Electron microscopy, biochemistry and pigment analysis and part of the biophysical analysis.

CM collected and analysed the data from DUAL-PAM,and Joliot spectroscopy, and measured the 77K fluorescence spectra of thylakoids). DAS performed single-particle negative staining electron microscopy (EM) experiments, including data collection, image analysis, and interpretation, and contributed to result discussions. AFB measured the time-resolved fluorescence kinetics on thylakoids and performed preliminary data analysis. EE measured time-resolved kinetics on the purified PSI complexes, and performed the global and target analysis of all time-resolved data sets. RC supervised the work in Amsterdam and contributed to the analysis and interpretation of the data. MS trained QCF on protein purification and purified the PSI and the L-PSI-LHCII supercomplex. RL performed Mass spectroscopy experiments (data collection and analysis). DG and colleagues organized and performed all sample collections.

QCF prepared most figures and additional files with the contribution of CM and EE.

CJ and QCF wrote the manuscript, with contributions from EE and RC. DAS, MS, DG, RC, RL edited and reviewed the manuscript. All authors read and approved the manuscript.

## Acknowledgements

We are grateful to Sandrine Ruitton, Pascal Mirleau, Deny Malengros, Fabrice Garcia, Michel Lafont, Sandrine Chenesseau, Frédérique Legendre and Bruno Belloni from the OSU Institut Pytheas (Aix Marseille University, CNRS, IRD, IRSTEA, OSU Institut Pythéas, Marseille, France) for the *Posidonia oceanica* samples collection. *Z. marina* and *Z. Noltii* were gifted by Pascal Bazile (8 Vie Pour la Planète, Projet Zorro) and *C. nodosa* by Heike Molinaar (Université Côte d’Azur).

The electron microscopy experiments were performed on the PiCSL-FBI core facility (Nicolas BROUILLY, Fabrice RICHARD and Aïcha AOUANE, IBDM, AMU-Marseille), member of the France-BioImaging national research infrastructure (ANR-10-INBS-04). A special thanks to Aïcha Aouane for her expertise and help in the plant samples preparation for electron microscopy. Thanks to Cécile Lecampion for the statistics analysis on the grana size quantification. We thank Dr. Kastritis and Dr. Hamdi for their assistance with transmission electron microscopy at Martin Luther University Halle-Wittenberg, Germany, and appreciate their support in providing access to the microscopy facility. Thanks to Rainer Hienerwadel for his expertise with the Joliot spectroscopy for ETR measurements and to Alexis Riche for his help and advice for 77 K fluorescence experiments. Thanks to Stefano Caffarri for the correction of an early version of the manuscript and general advice on biochemistry techniques. We are grateful to EMBO who granted QCF with a Scientific Exchange Grant (number 9348) and thanks to Alexey Amunts. Huge thanks to Laurie Casalot, Lea Sylvi, Manon Bartoli, Corinne Valette and Sophie Guasco for providing the equipment necessary for sample preparation. We are deeply grateful to Paul Hudson for providing a scientific environment of excellence.

