## Supplementary material for "Thriving Across Depths: How Blue Light Shapes a Large PSI Supercomplex and Specific Photosynthetic Traits in the seagrass *Posidonia oceanica*": Suplemental Figure

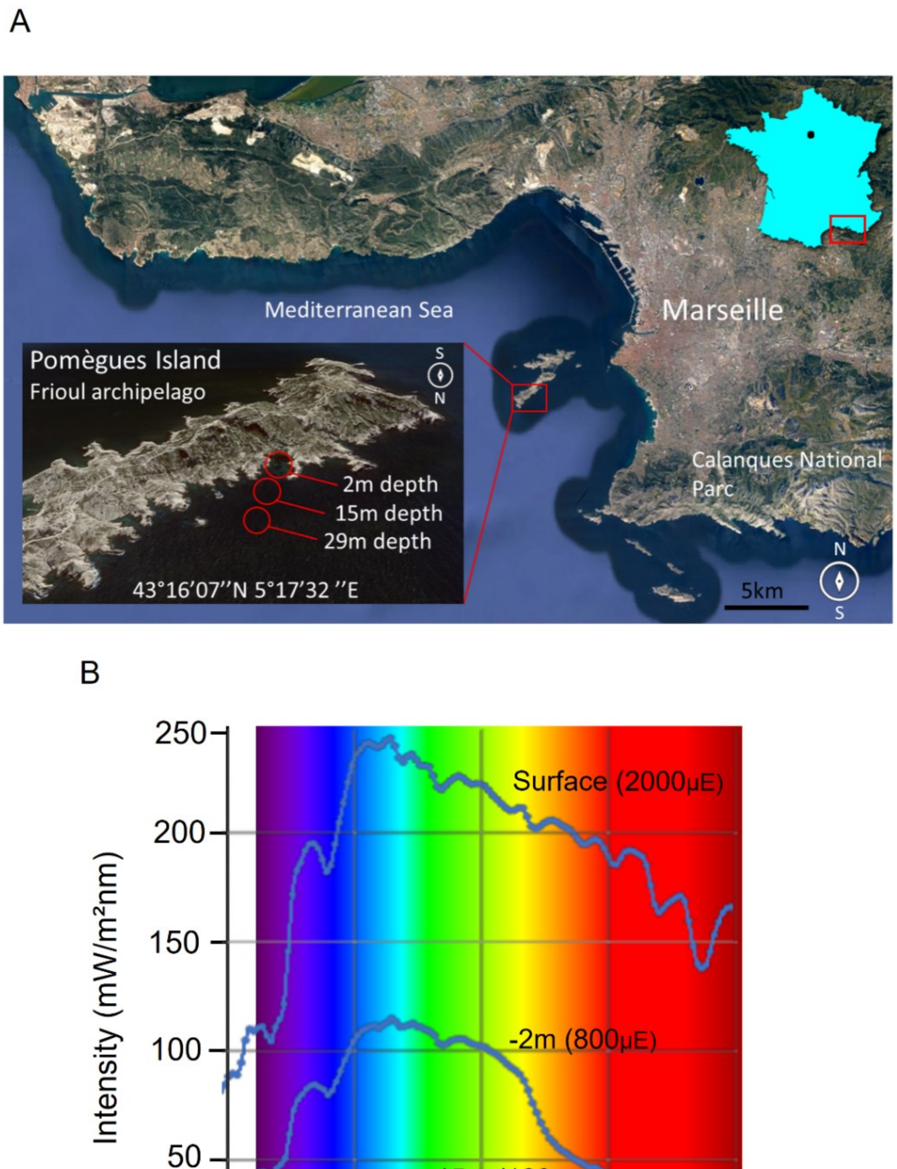


**Figure S1: Geographic localization and the irradiance reaching the selected *P. oceanica* meadows.** **(A)** The *P.oceanica* plants used for this study were harvested from 2, 15, and 26m deep meadows located in the Frioul Archipelago, a Mediterranean island territory of Marseille municipality (south of France). The harvesting site is 20 minutes from the Marseille harbor by boat. The meadows of interest grew at a 2, 15, and 26m depth and belonged to a higher order discontinuous patch covering the coastal border of the Pomègues island. Right after harvest, the shoots were stored in the dark in an isolated box filled with ice and seawater. **(B)** The light spectrum was measured in March 2019 and irradiance is expressed in µmol photons.m²/s (µE).

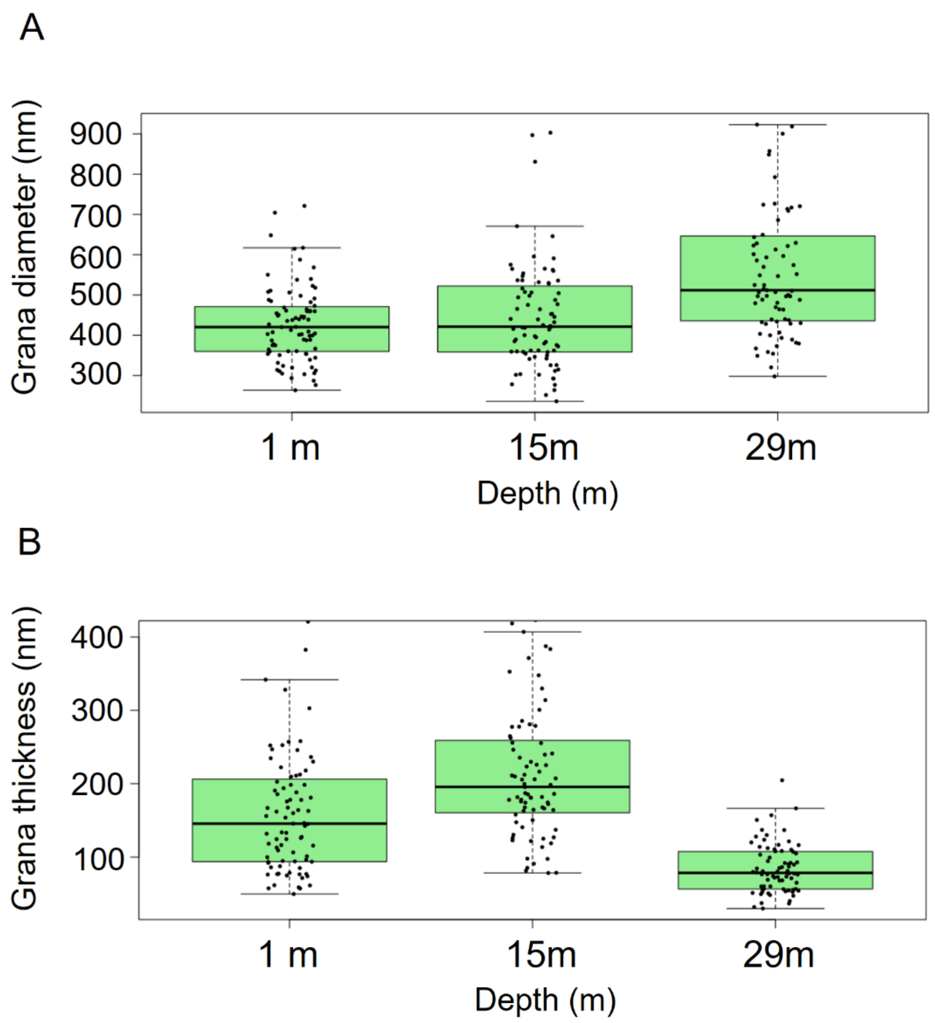


**Figure S2: Distribution of the diameter and thickness of thylakoids grana at different growing depths (2, 15, and 26m deep).**

The diameter **(A)** and thickness **(B)** of the grana were measured from electron micrographs using ImageJ software. The pixel length of the selected area was normalized by the scale bar length of each used micrograph (around 10 for each depth). Around 80 grana were selected for each tested depth and each dot from the dot-plot represented one measurement. Blue boxes contain 50% of the measurement and the black line is the median.

A

| Chl *a/b* | 2m depth | 15m depth | 26m depth |
|---|---|---|---|
| **Winter 2019** | 2.21 ± 0.01 | 2.05 ± 0.04 | 2.02 ± 0.02 |
| **Summer 2020** | 1.92 ± 0.02 | 1.93 ± 0.01 | 1.91 ± 0.01 |
| **Winter 2021** | 2.13 ± 0.04 | 2 ± 0.04 | 2.01 ± 0.02 |
| **Summer 2021** | 1.85 ± 0.01 | 1.83 ± 0,01 | 1.82 ± 0.03 |
| **Winter 2022** | 2.14 ± 0.01 | 2.04 ± 0.01 | 2.06 ± 0.02 |

B

| Total LHCII/PSII+PSI | 2m depth | 15m depth | 26m depth |
|---|---|---|---|
| **Winter 2019** | 6.4 | 7.5 | 8,2 |

C

| *A.thaliana* | HL | NL | LL |
|---|---|---|---|
| **Chl *a / b*** | 3.2 | 3 | 2.7 |
| **Total LHCII/PSII+PSI** | 3.2 | 3.8 | 5.1 |

**Figure S3: Regulation of the LHCII in *P. oceanica* at different seasons and years of collection.**
**(A)** Chlorophyll *a*/*b* stoichiometry of the purified thylakoids from *P. oceanica* leaves collected at 2, 15, and 26m depth over different years and seasons. Data are expressed as the mean ± SD, n = 3 for Winter 2019, 2021 and summer 2020, and n = 2 for summer 2021 and winter 2022. In summer, the Chl *a*/*b* is lower, yet no significant variation is measured between tested depths. **(B)** Estimation of the LHCII number per PSII + PSI reaction center based on the Chl *a*/b from purified thylakoids in winter 2019 and the Chl content of the isolated LHCII and PSI as well as the PSI-PSII ratio from winter 2019. The Chl content of PSII was taken from Su et al,. 2017(Su et al., 2017). The estimation was performed as in Ünlü et al., 2014(Ünlü et al., 2014), using the PSI/PSII ratio, the PSI-LHCI and L-PSI-LHCII relative content, the Chl *a/b* of purified thylakoids, and the Chl content of isolated LHCII and PSI-LHCI complexes **(C)** The same procedure as in (B) was applied to *A.thaliana* based on the Chl *a/b* from literature(Kouřil et al., 2013)

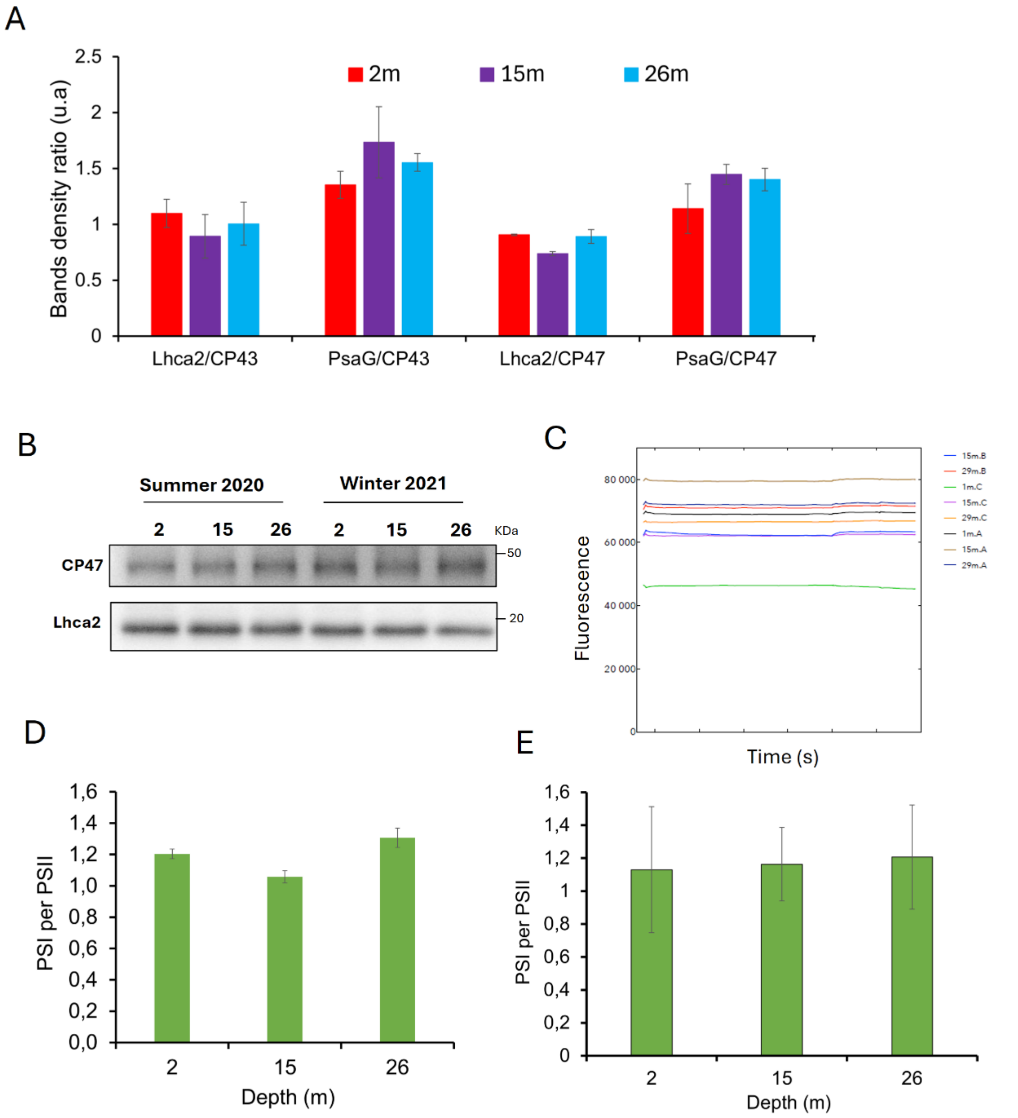


**Figure S4: Immunodetection of the photosystems stoichiometry and phosphorylation of the LHCII trimers in *P. oceanica* at different growing depths, seasons, and years.** **(A)** The immunoblot band pixel density (**Figure 2B**) was quantified using the ImageJ software. The band ratio is expressed by dividing the pixels of PSI subunits by the pixel density of PSII subunits. The data are expressed as the mean ± SD, n = 2. Note: the value of the ratio is not representative of the absolute PSI/PSII stoichiometry. **(B)** Evaluation of the PSI/PSII stoichiometry by immunoblot analysis. The thylakoid membranes were extracted from *P. oceanica* collected at 2, 15, and 26m depth at different seasons and years. 0,5µg of Chl was loaded on the gel. Immunodetection was performed using specific primary antibodies (Agrisera) as indicated. **(C)** Fluorescence kinetics are measured by ECS to control the DCMU+HA inhibition of photosystem II for the ECS estimation of PSI/PSII stoichiometry. **(D-E)** The PSI/PSII stoichiometry was estimated from fresh young leaves in winter based on the difference of ECS signal between untreated (PSII+PSI) and DCMU+HA-infiltrated (PSI only) leaves. Data is expressed as the mean from two independent experiments performed two weeks apart: January 2019 (d) and February 2019 (e) at the 3 tested depths, The PSI/PSII ratios are plotted as mean ± SD, n = 3.

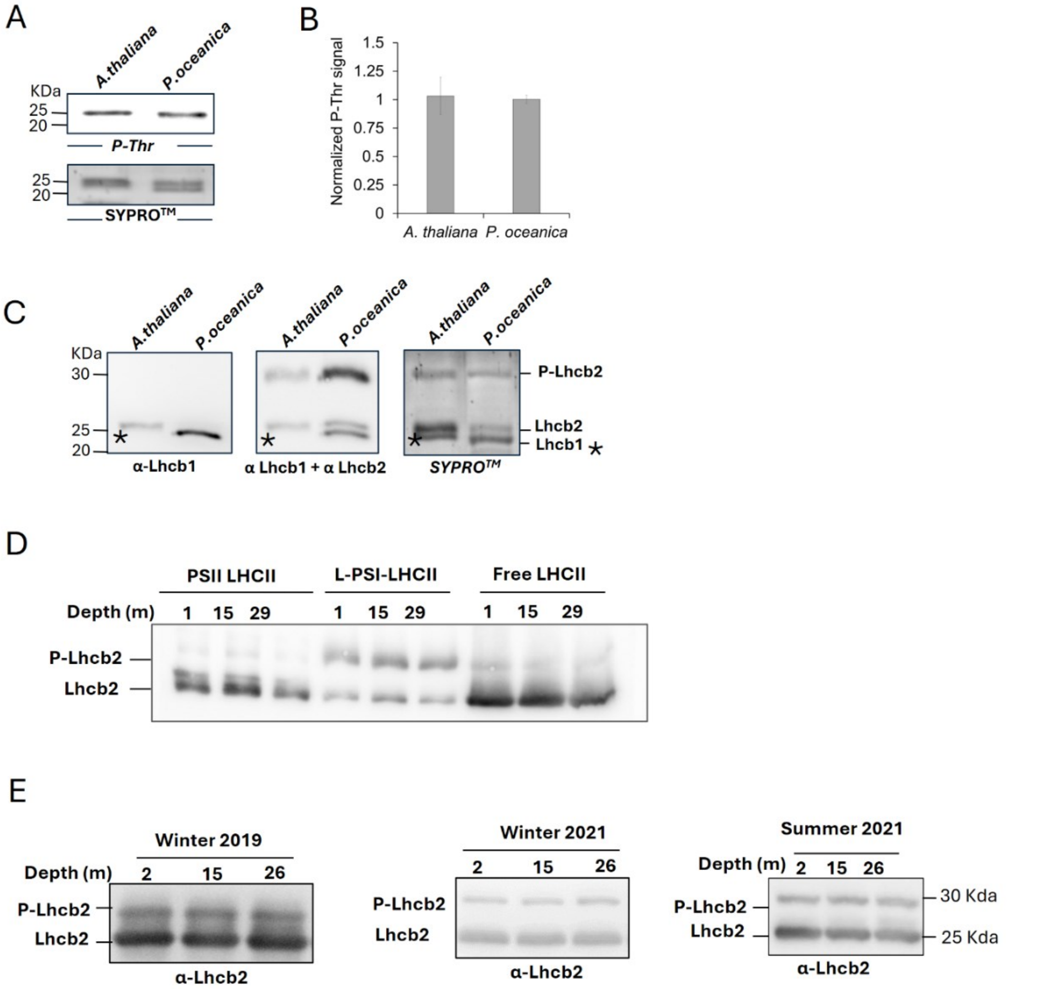


**Figure S5: Analysis of the bound and free LHCII phosphorylation in *P. oceanica*.** **(A-C)** Second-dimension analysis of the bound LHCII of the PSI-LHCII from *A. thaliana* and the L-PSI-LHCII from *P. oceanica.* Complexes were isolated in the first dimension by BN-PAGE. The bands were sliced and deposited on the top of a 13% acrylamide/bis-acrylamide gel. **(A)** Top: Immunodetection was performed using an anti-Phospho-threonine antibody (Cell Signaling). Bottom: A second gel was performed and stained with SyproRuby. **(B)** The band's pixel density from (A) was quantified using ImageJ software. The density from the P-Thr immunodetection bands was normalized against the signal from the SyproRuby stained bands (corresponding to Lhcb1 and Lhcb2). The data is expressed as the mean ± SD, n = 3. **(C)** Phos-tag 2D-PAGE Immunoblot using Lhcb1 and Lhcb2 antibodies (Agrisera). The same membrane was used for the two blots. A second gel was performed but stained with Sypro Ruby. **(D)** Phos-tag 2D-PAGE Immunoblot using Lhcb2 antibodies (Agrisera). For the 3 tested depths, the bands corresponding to PSII-LHCII, L-PSI-LHCII and free LHCII from 1D Native-PAGE were cut, denatured and loaded on the top of SDS denaturing gel. **(E)** Phosphorylation of the LHCII in *P. oceanica* thylakoids from 2, 15, and 26m depths collected during different seasons and years. 0.5µg of chlorophyll was loaded on a 13% acrylamide/bis-acrylamide gel supplemented with 50µM Phos-tag$^{TM}$ + 100µM MnCl2. Immunodetection was performed using specific primary antibodies (Agrisera) as indicated.

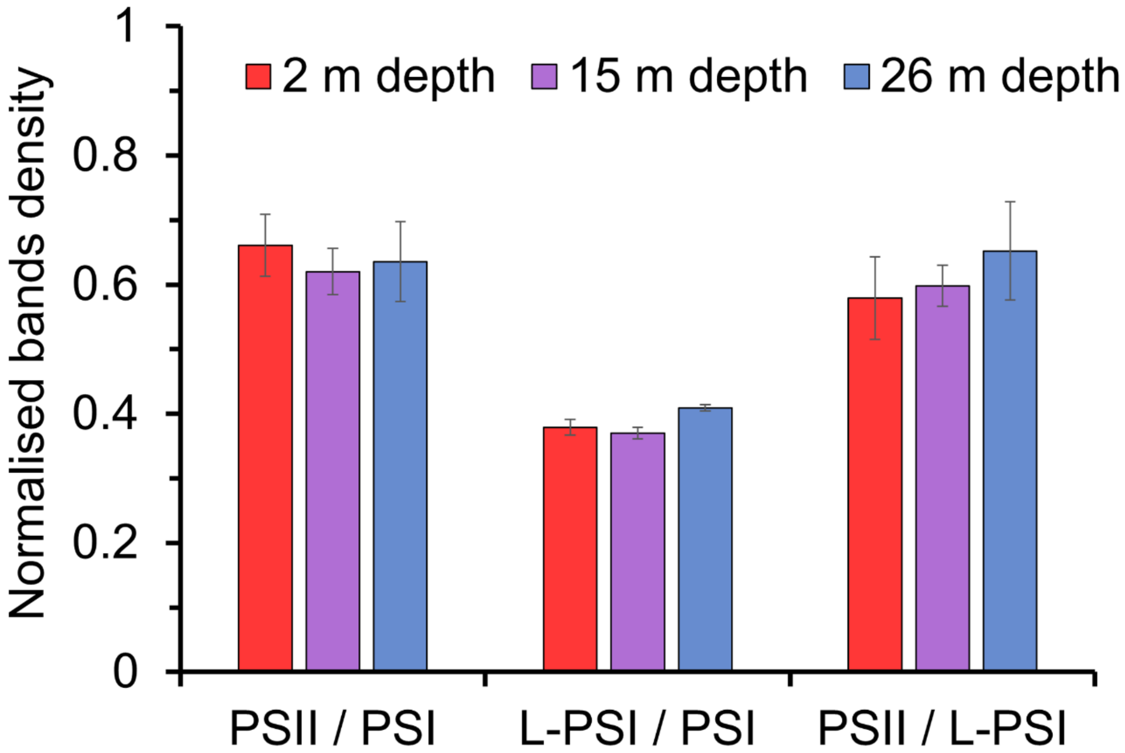


**Figure S6: Regulation of the L-PSI-LHCII and other photosystem supercomplexes between the three-tested depth.** The pixel density of the bands from **Fig. 3A** was quantified using ImageJ software, and the normalized intensity was plotted. The data are expressed as the mean ± SD, n = 3.

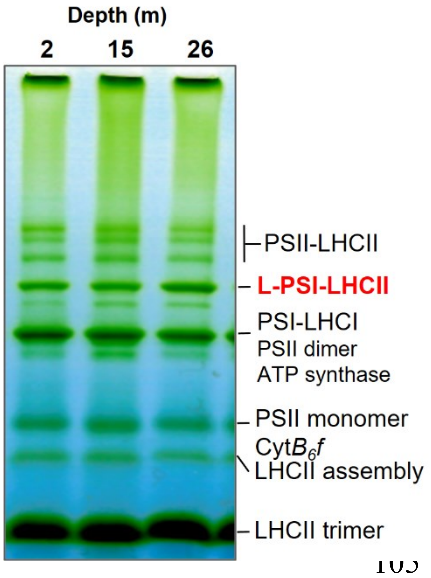


**Figure S7: Regulation of the L-PSI-LHCII and other photosystem supercomplexes between the three tested depths in summer.**

BN-PAGE of solubilized thylakoid membrane proteins extracted from *P.* oceanica at different growing depths and collected in summer. About 5µg of Chl was loaded onto the gel.

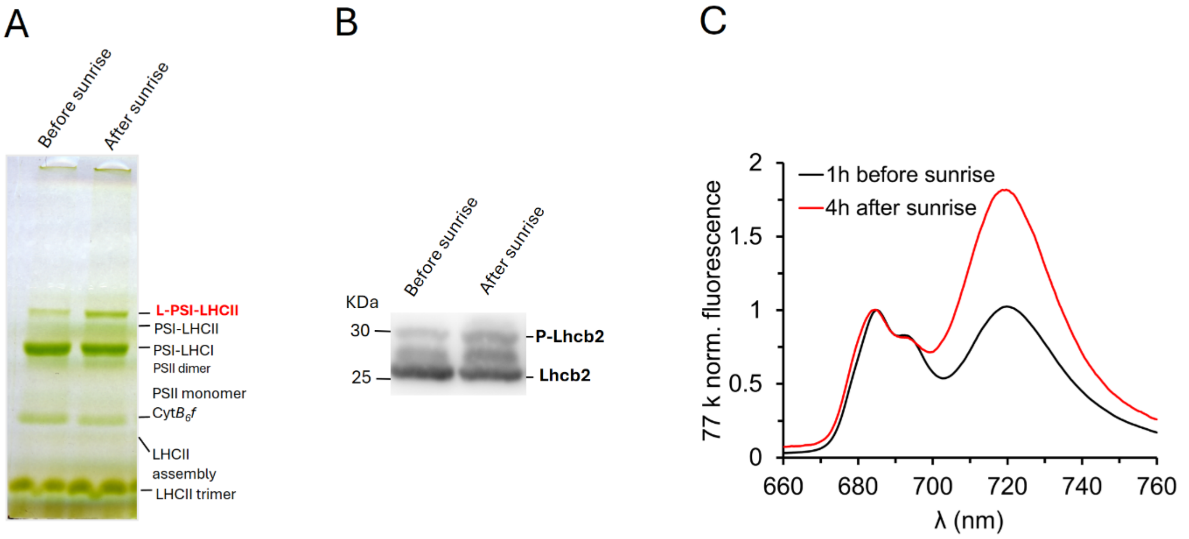


**Figure S8: Induction of the L-PSI-LHCII by sunlight.**
**(A)** CN-PAGE of solubilized thylakoid membrane proteins extracted from *P.* oceanica collected at a depth of 15m before and after sunrise *in situ*. For each tested depth, about 5µg of chlorophyll was loaded onto the gel. **(B)** Phos-tag immunoblot targeted against Lhcb2 on thylakoids (0.5 µg of Chl were loaded). **(C)** 77k emission fluorescence of thylakoid membranes.

| | ***P. o*** **PSI-LHCI** | ***P. o*** **L-PSI-LHCII** |
|---|---|---|
| **chl *a*/chl *b* in acteone** | 6.84 ±0.05 | 4.33 ±0.03 |
| **Predicted chl *a*+chl *b*** | 157 | 227 |
| **Estimated chl *a*** | 137 | 184,5 |
| **Estimated chl *b*** | 20 | 42,5 |

**Figure S9: Estimation of the chlorophyll content of the PSI-LHCI and the L-PSI-LHCII from *P.oceanica*.**
Chlorophyll content (extracted in acetone) of the purified PSI-LHCI and L-PSI-LHCII from *P. oceanica* and the PSI-LHCI from *A. thaliana*. The total number of chlorophylls (predicted Chl *a* + Chl *b*) for *P. oceanica* was based on the total chlorophyll number in the PSI-LHCI from the PSI-LHCI-LHCII model *A. thaliana* (Wu et al., 2023). For *P. oceanica* PSI-LHCII, the number of chlorophylls was the sum of the chlorophyll numbers estimated for the *P. oceanica* PSI-LHCI, and LHCII trimer and the Lhca1-Lhca4 dimer (estimated from the number in *A.thaliana* structure).

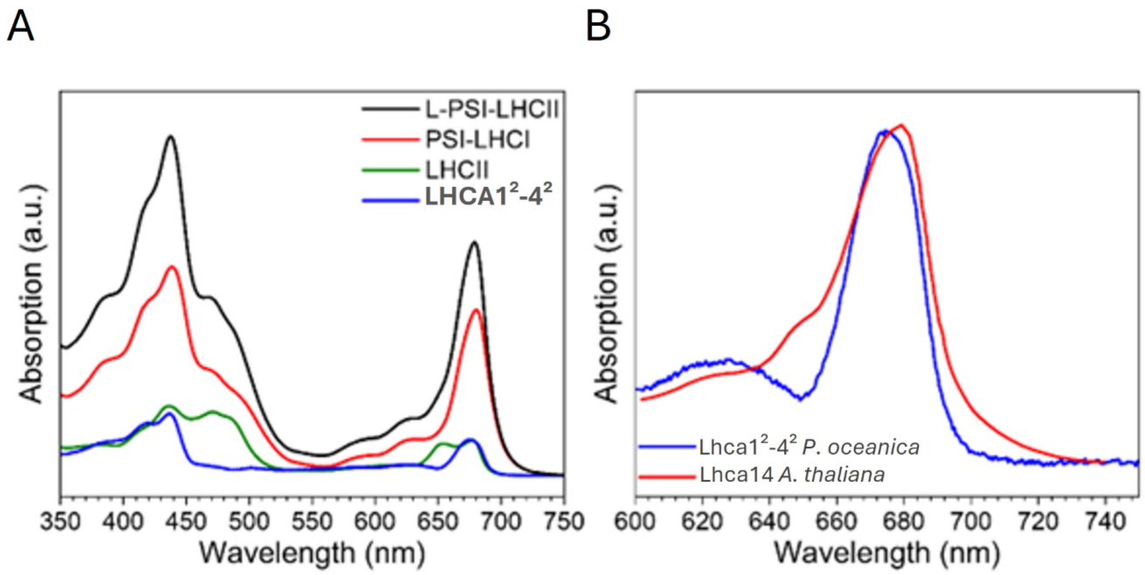


**Figure S10: Absorption spectra determined from the spectral deconvolution and comparison of the Lhca14 absorption spectrum. (A)** Absorption spectra as determined for the L-PSI-LHCII, PSI-LHCI, LHCII trimer and Lhca1$^2$-Lhca4$^2$ (Lhca1$^2$-4$^2$) complex by scaling to their Chl content (see method). **(B)** Comparison of the xLhca14 absorption spectrum from the spectral deconvolution of *P. oceanica* to the Lhca1-4 absorption spectrum of *A. thaliana* (Wientjes et al., 2011).

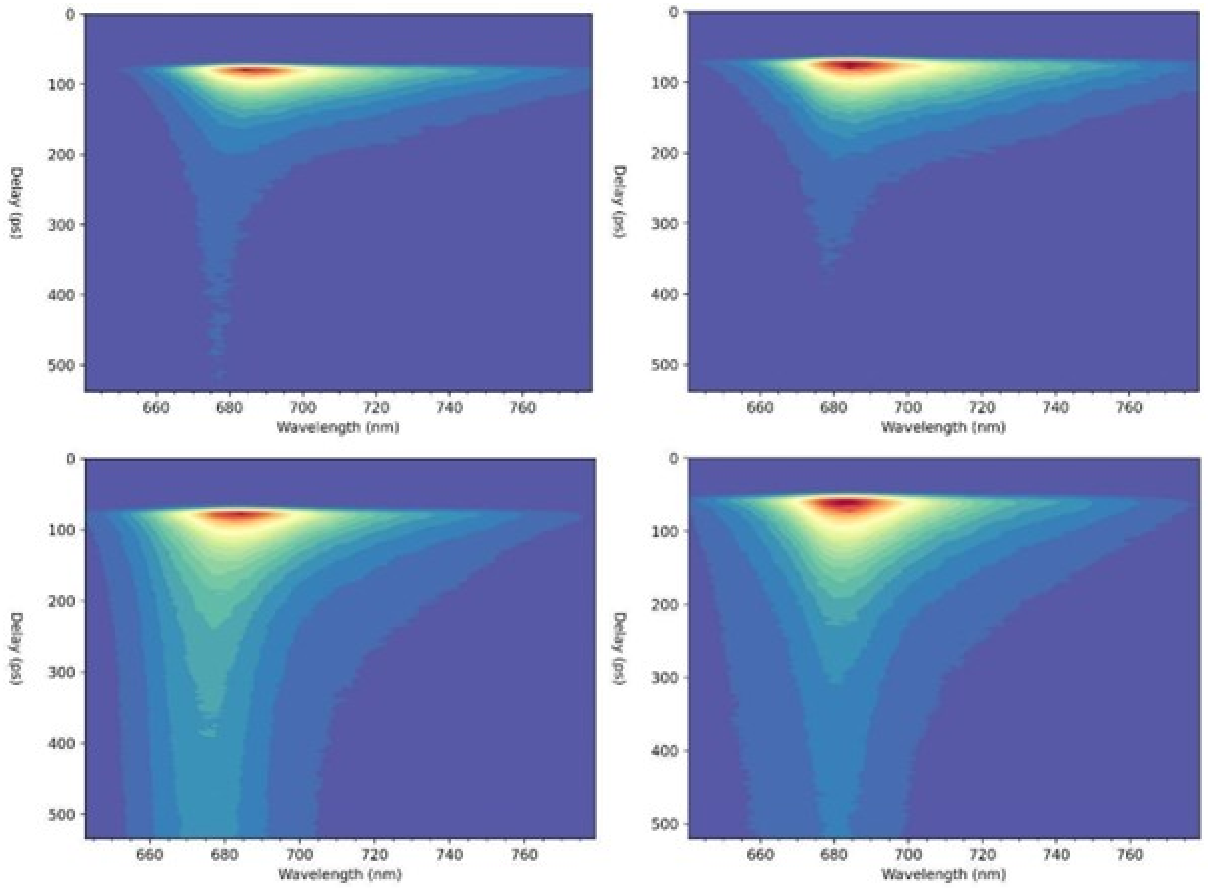


**Figure S11: PSI-LHCI and L-PSI-LHCII streak camera images.** Top-left: PSI-LHCI excited at 400 nm. Top-right: PSI-LHCI excited at 475 nm. Bottom-left: L-PSI-LHCII excited at 400 nm. Bottom-right: L-PSI-LHCII excited at 475 nm.

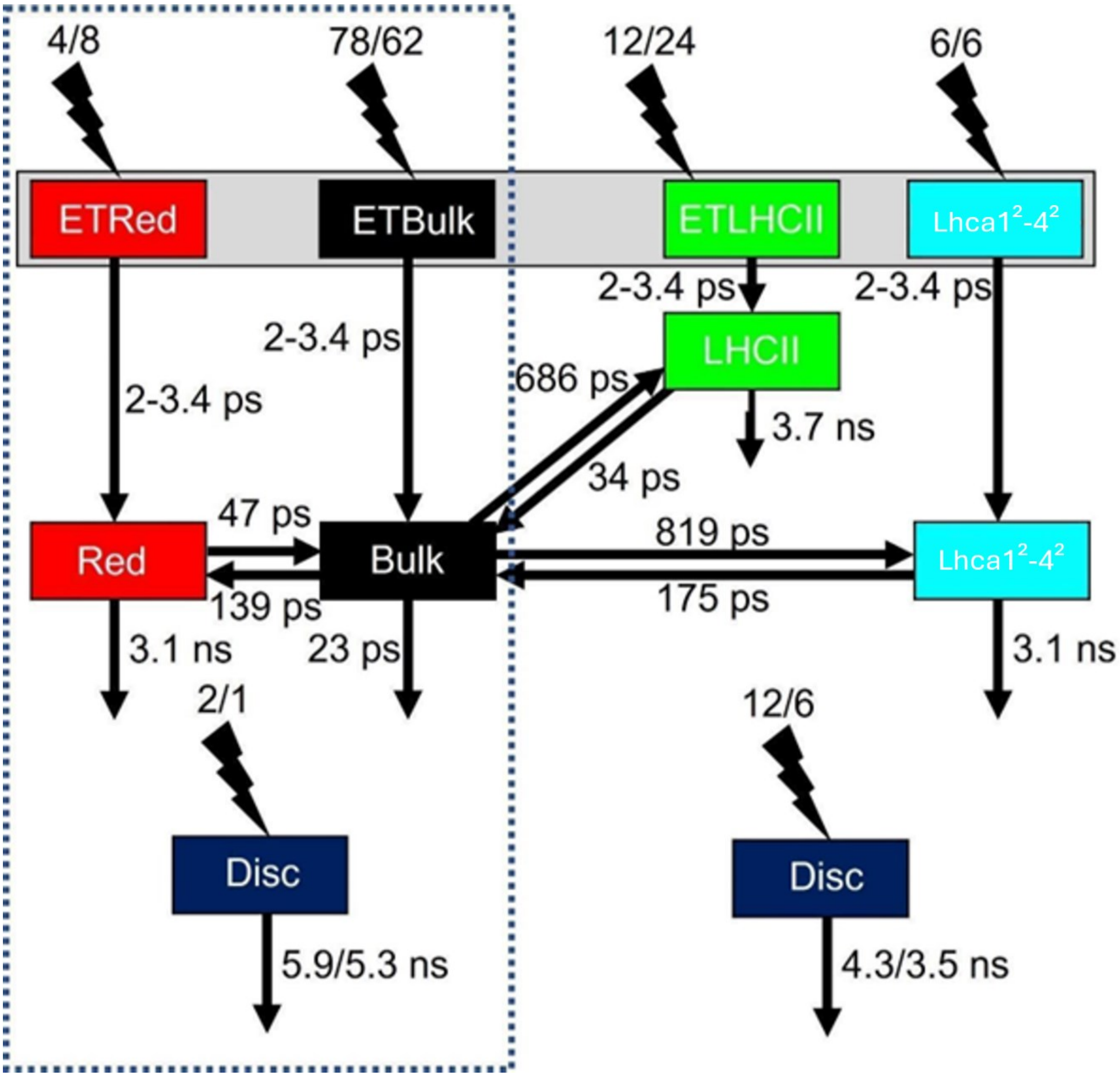


**Figure S12: Full target kinetic scheme of the L-PSI-LHCII.** The relative initial excitation densities are shown in percentages above the lightning symbols. The left number indicates the relative excitation density for the 400 nm experiments and the right number for the 475 nm ones. The PSI-LHCI kinetic model is marked by the blue dashed rectangle. The Lhca$1^2$-Lhca$4^2$ dimer is referred to as Lhca$1^2$-$4^2$.

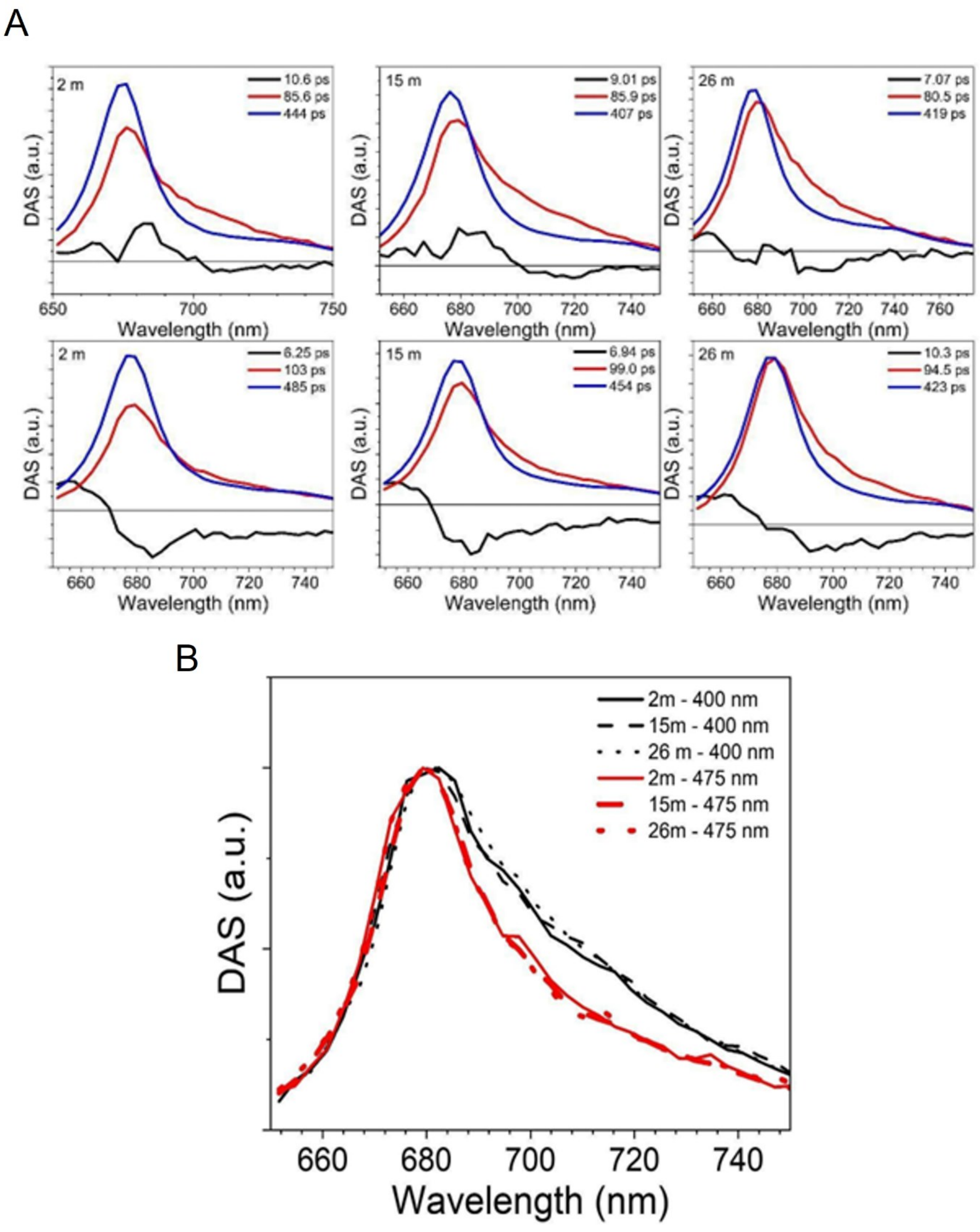


**Figure S13: Comparison of TRF spectra at different depths in summer and winter.** **(A)** Decay-associated spectra obtained from global fitting of time-resolved fluorescence spectra. Thylakoids from leaves collected at depths of 2, 15, and 26m in summer were excited at 400 (top) and 475 nm (bottom). **(B)** Normalized PSI DAS overlap at different depths for winter thylakoids.

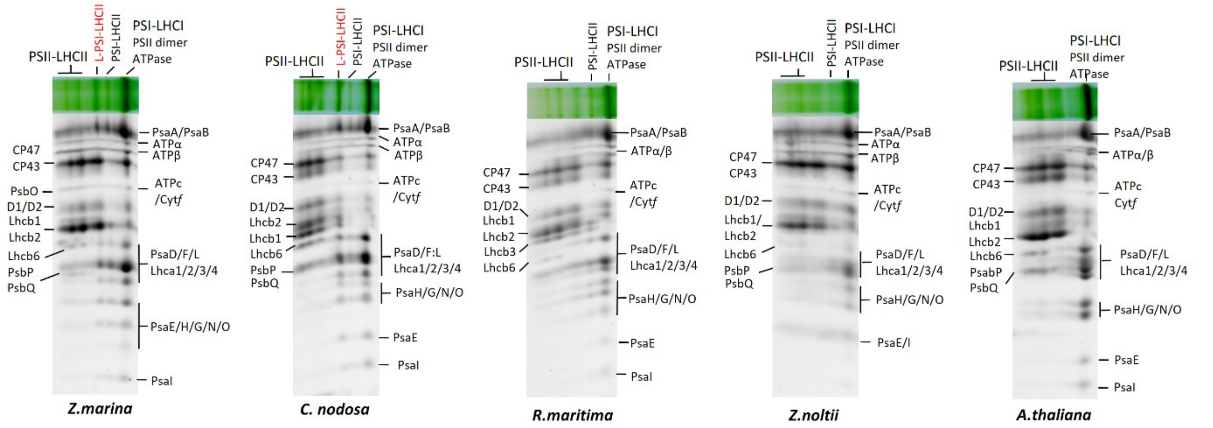


**Figure S14: Identification of the peptide content of BN-PAGE bands by a second dimension on Urea-PAGE.** The region of the BN-PAGE from Figure 8 containing PS's supercomplexes was sliced, denatured and deposited on top of a 13% acrylamide-bis acrylamide gel. Gel stained using SyproRuby.

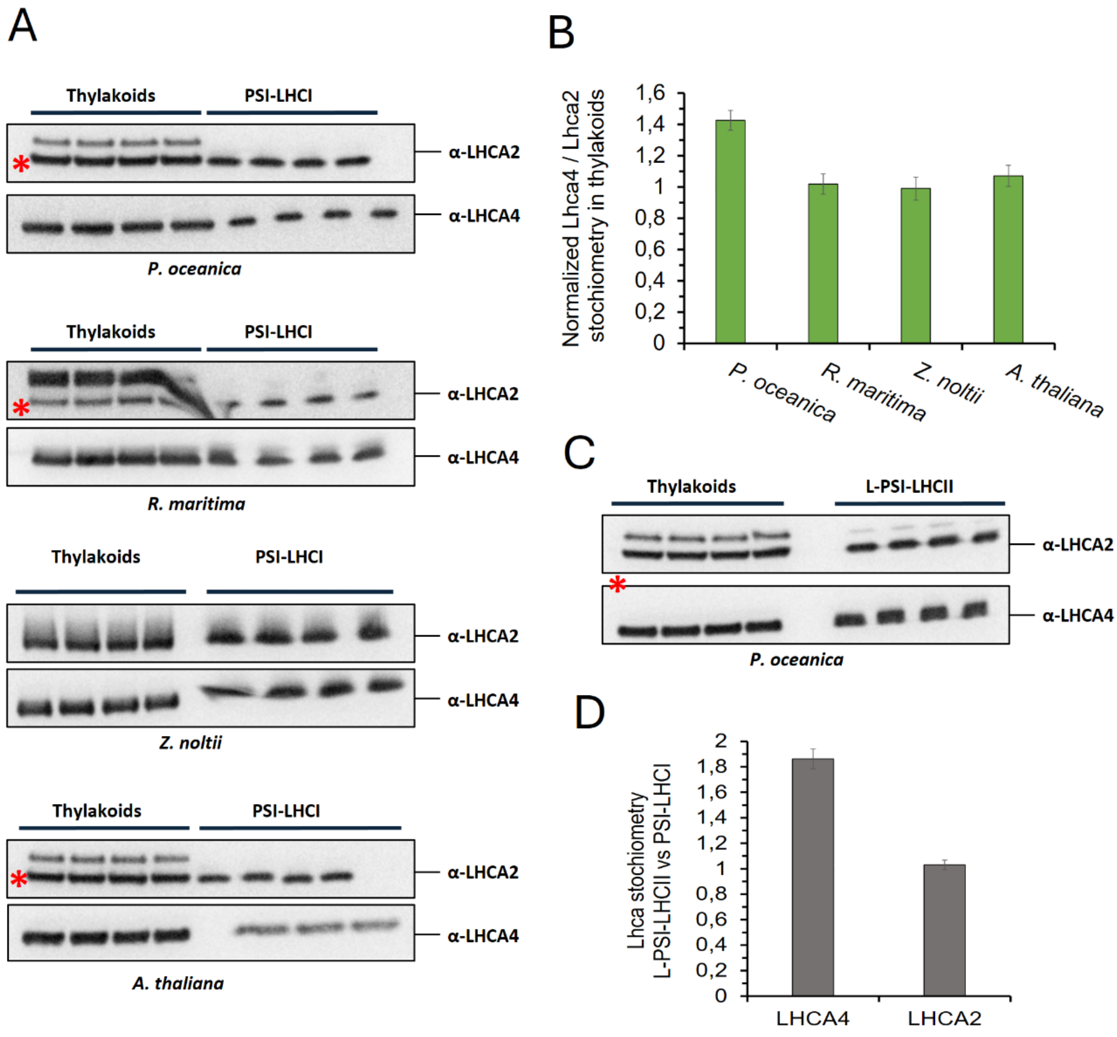


**Figure S15: Lhca4 stoichiometry in the thylakoids of *P. oceanica* compared to shallow growing seagrasses and *A. thaliana*.**
**(A)** Immunodetection of Lhca4 and Lhca2 protein from thylakoids and the isolated PSI from *P. oceanica*, *R. maritima*, *Z. noltii* and *A. thaliana*. For the thylakoids, 0.5 µg of chlorophyll was loaded on the top of the gel. L-PSI-LHCII from *P. oceanica* and the PSI-LHCI from indicated species were isolated by BN-PAGE. After separation, the bands were sliced, denatured and deposited on the top of the same gels together with the thylakoids extracts. For Lhca2 blot, the red asterisk indicates the band used for quantification. **(B)** Quantification of Lhca2 and Lhca4 protein stoichiometry in the thylakoids of *P. oceanica*, *R. maritima*, *Z. noltii* and *A. thaliana*. Corresponding immunoblots are presented in the (A). The Lhca4/Lhca2 stoichiometry was determined by normalizing their signal in thylakoids to the signal from PSI-LHCI isolated in each species. Data are presented as the mean ± SD, N=4. **(C) :** Immunodetection of Lhca4 and Lhca2 protein from thylakoids and the isolated PSI from *P. oceanica*. **c :** Estimation of the stoichiometry of Lhca2 and Lhca4 from the isolated PSI-LHCI and L-PSI-LHCII in *P. oceanica*. For both Lhca2 and Lhca4 proteins, the pixel density of the bands from PSI-LHCI and the L-PSI-LHCII was normalized by the pixel density of the bands from their respective thylakoids fraction. Data are presented as the mean ± SD, N=4.

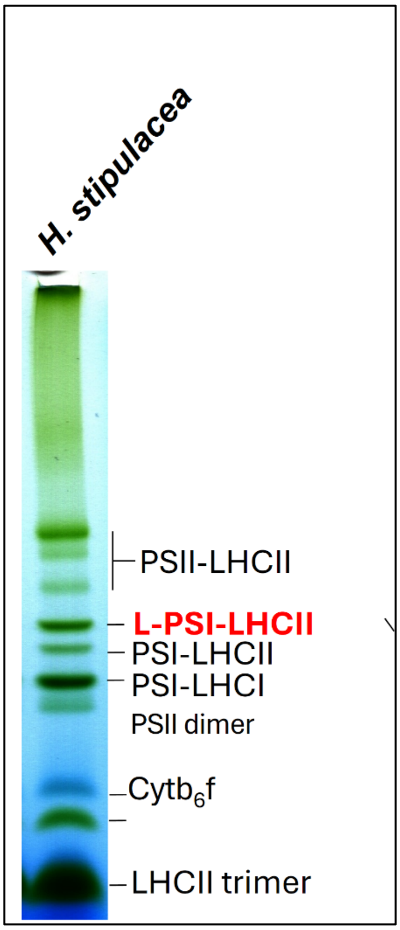


**Figure S16: Photosystem supercomplexes in *Halophila stipulacea***

BN-PAGE of solubilized thylakoid membrane proteins extracted from *H. stipulacea*. About 5µg of Chl was loaded onto the gel.

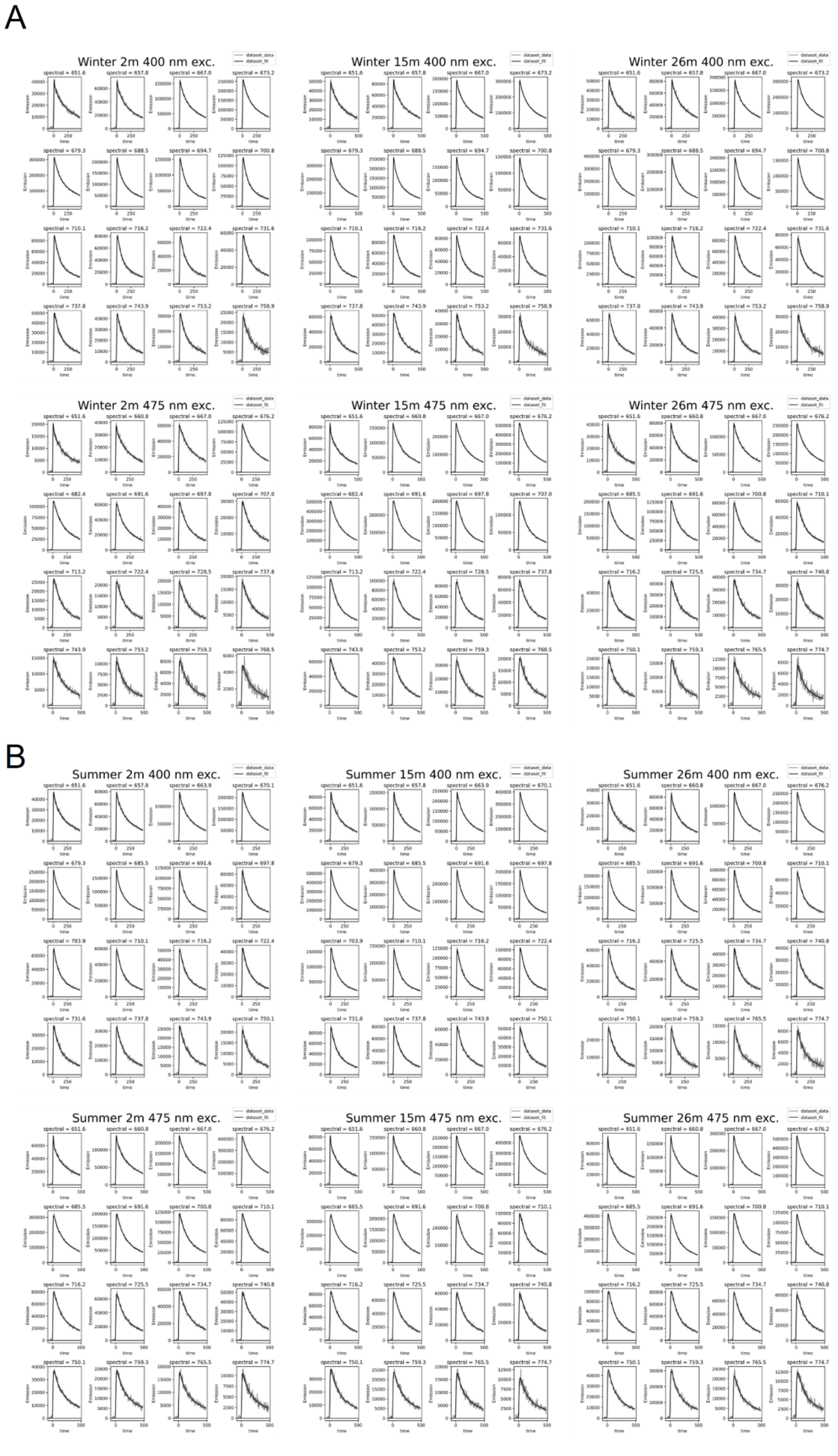


**Figure S17: Fitting results of the fluorescence traces from the target analysis.** (a) Winter, (b) summer.

### Extended data references :


Kouřil, R., Wientjes, E., Bultema, J. B., Croce, R., and Boekema, E. J. (2013). High-light vs. low-light: Effect of light acclimation on photosystem II composition and organization in Arabidopsis thaliana. *Biochimica et Biophysica Acta (BBA) - Bioenergetics* 1827:411–419.

Su, X., Ma, J., Wei, X., Cao, P., Zhu, D., Chang, W., Liu, Z., Zhang, X., and Li, M. (2017). Structure and assembly mechanism of plant C2S2M2-type PSII-LHCII supercomplex. *Science* Advance Access published August 25, 2017, doi:10.1126/science.aan0327.

Ünlü, C., Drop, B., Croce, R., and van Amerongen, H. (2014). State transitions in *Chlamydomonas reinhardtii* strongly modulate the functional size of photosystem II but not of photosystem I. *Proc. Natl. Acad. Sci. U.S.A.* 111:3460–3465.

Wientjes, E., van Stokkum, I. H. M., van Amerongen, H., and Croce, R. (2011). The Role of the Individual Lhcas in Photosystem I Excitation Energy Trapping. *Biophysical Journal* 101:745–754.

Wu, J., Chen, S., Wang, C., Lin, W., Huang, C., Fan, C., Han, D., Lu, D., Xu, X., Sui, S., et al. (2023). Regulatory dynamics of the higher-plant PSI–LHCI supercomplex during state transitions. *Molecular Plant* 16:1937–1950.
